# Dissecting the role of PCNA and Pif1 in replication of individual DNA molecules by DNA polymerase δ

**DOI:** 10.64898/2026.05.19.725978

**Authors:** Manal S. Zaher, Carrie Stith, Ayush Mistry, Purna B. Poudel, Edwin Antony, Peter M. Burgers, Roberto Galletto

## Abstract

PCNA loading and stabilization at primer-template junctions is crucial for processive DNA synthesis by replicative polymerases. It is also essential for strand displacement synthesis during Break Induced Replication (BIR). In this study, we employed a single molecule approach to directly visualize the role of PCNA, DNA polymerase δ and the Pif1 DNA helicase in these pathways. We first dissected the preferential loading of PCNA by RFC to 3’-junctions of gaps, nicks and flapped substrates. Moreover, we found that Pol δ and Pif1 both stabilize loaded PCNA and reduce its diffusion coefficient along DNA. We further show that the interaction of PCNA with Pol δ is essential for gap-filling synthesis. On a 5’-flapped substrate, we show that both Pol δ and Pif1 are required for strand displacement synthesis, while PCNA is not; although, it stimulates the process.

## Introduction

DNA replication occurs via a highly orchestrated series of events that lead to DNA synthesis by the replicative DNA polymerases ε and δ of the leading and lagging strands, respectively (1-3). During lagging strand DNA replication, one of the central steps in the process is loading of Proliferating Cell Nuclear Antigen (PCNA) sliding clamp onto DNA by Replication Factor C (RFC) (4-6). Recruitment and stabilization of PCNA at the 3’-end of a primer template junction is necessary for interaction with Pol δ and formation of a complex competent for DNA synthesis. Over the past 40 years, ensemble and structural studies of this step have provided a wealth of information (5, 7-10) that is now being expanded by the ability to image this step at individual DNA sites (11-15). Formation of a PCNA-Pol δ complex leads to processive DNA synthesis, filling the gaps left behind by the preceding Okazaki fragment. In this process, the PCNA-Pol δ complex must be able to displace the single-stranded (ss)DNA binding protein (RPA) tightly bound to the template strand, and complete DNA synthesis of the gap up to the start of the RNA primer previously deposited by the activity of the Pol α-primase complex. Upon completion of gap filling, Pol δ displaces the downstream RNA via strand displacement DNA synthesis allowing the flap endonuclease 1 (FEN1) to process it. This generates a ligatable nick that is then sealed by DNA ligase I (16-19). Failure to organize these activities can lead to the accumulation of toxic DNA intermediates (20-24).

Importantly, the strand displacement activity of Pol δ must be limited to formation of 5’-flaps shorter than ∼10 nt, as longer ones can be bound by RPA thereby suppressing the activity of FEN1 (18, 25, 26). During proper Okazaki fragment maturation, the ∼10 mer RNA primer is displaced by Pol δ, but this RNA cannot be used as an entry point by the Pif1 helicase to elicit unwinding of the downstream duplex (27). Furthermore, strand displacement synthesis by Pol δ is generally coupled to degradation of the displaced strand by FEN1 (28). Therefore, involvement of the Pif1 helicase during DNA replication is not expected to occur, unless the principal pathway goes awry, and longer stretches of DNA are being displaced and ssDNA exposed. In this eventuality, a second pathway that uses the Dna2 helicase/nuclease and the Pif1 helicase must be activated for flap processing (29, 30). In this regard, we showed that even a small amount of ssDNA transiently generated by Pol δ via strand displacement can be an entry point for the Pif1 helicase (31). However, these events must be tightly suppressed as they would lead to extensive DNA re-replication into the previously synthesized Okazaki fragment.

The situation is very different during Break Induced Replication (BIR) where strand displacement synthesis must occur efficiently for the process to proceed. BIR is a last resort recombination-dependent DNA pathway to repair broken DNA ends. During BIR, the 3’-ssDNA of a broken end coated by the recombinase Rad51 invades the sister chromatid and pairs with its homologous sequence to form a D-loop (32, 33). At this stage, PCNA loading is required to form a processive complex with Pol δ to allow copying kbps of information from the sister chromatid, via strand displacement DNA synthesis, into regions that are bound by proteins and wrapped into nucleosomes (34, 35). However, neither Pol δ nor PCNA-Pol δ complexes can carry out substantial amounts of strand displacement DNA synthesis on their own. Indeed, efficient BIR in yeast and human cells depends on the activity of the Pif1 helicase (36-39), via its interaction with PCNA (36, 37). The large amount of DNA synthesized during BIR points to DNA unwinding by Pif1 as being the crucial step in the process that provides the activity needed to open the downstream duplex and expose the template to be copied by Pol δ. Consistent with this idea, we showed that *in vitro* Pif1 stimulates strand displacement DNA synthesis by PCNA-Pol δ, even when the downstream duplex is bound by an array of dsDNA binding proteins or wrapped into a nucleosome (31, 40). We also showed that the helicase activity of Pif1 or Pfh1 is dominated by a highly processive cycle of DNA unwinding and rewinding that effectively limits its activity to the opening of only short stretches of DNA, no longer than 20 bp (41, 42). Therefore, stimulation of strand displacement DNA synthesis by Pif1 during BIR most likely arises from processive opening of short stretches of DNA by the 5’-3’ helicase activity of Pif1, coupled to concomitant DNA synthesis by Pol δ on the opposite strand. This mechanism would enable long-range DNA synthesis during BIR (34, 36, 38). It would also effectively couple DNA unwinding to DNA synthesis if stable G-quadruplexes stall the replication fork. Indeed, Pif1 helicases facilitate transient disruption of these stable G-quadruplex structures to allow DNA synthesis to progress (31, 43, 44).

Independent of the pathway we examine, PCNA loading onto DNA is the crucial step common to all (32, 36, 45). Indeed, PCNA is a central hub at the replication fork, with roles in DNA replication, repair, chromatin assembly and cell-cycle regulation (46). It acts as a sliding clamp that binds to replicative polymerases, Pol ε and Pol δ (4, 6, 47). Moreover, PCNA acts as scaffold that recruits upwards of hundreds of proteins to DNA through a conserved PCNA-interacting-protein (PIP)-box (46, 48, 49). These proteins are involved in a variety of DNA transactions during DNA replication, repair and recombination (46, 50). Therefore, failures in PCNA loading or turnover may lead to genome instability, replication stress and DNA damage (51, 52). PCNA is a ring-shaped homotrimer that topologically encircles and slides over double-stranded (ds)DNA (6, 15). As such, it needs a clamp loader to open its ring and load it onto DNA. This loading process is mediated by the heteropentameric AAA+ ATPase complex RFC (9, 10). RFC is arc-shaped and binds PCNA like a screw cap with a bottle (9). To load PCNA, RFC binds PCNA and opens its ring to form a stable complex which is ATP-dependent (8-10, 53). RFC then specifically recognizes and binds primer–template junctions containing a recessed 3′-OH end (5, 8). This DNA binding stimulates ATP hydrolysis which leads to the dissociation of RFC from loaded PCNA (7, 8). In addition, the N-terminal single BRCT domain of RFC1 binds the 5’-junction of gapped substrates, suggesting a role in gap repair (54, 55). Also, the BRCT domain of RFC1 is proposed to form a closed complex of RFC-PCNA-Pol δ that increases processivity of DNA synthesis during gap-filling (12). This is somewhat different from the behavior of a form of RFC, lacking this BRCT domain, which is more commonly used in biochemical studies (56).

Notwithstanding the importance of these pathways, studies that directly examine the coordination of the RFC-dependent loading of PCNA, Pol δ synthesis and Pif1 unwinding during gap-filling and strand displacement synthesis are limited. In this work, we employed optical trapping combined with confocal imaging to investigate the loading of PCNA by RFC onto individual DNA substrates with an available 3’-junction that can be used for DNA synthesis by Pol δ. We show that PCNA is preferentially loaded onto the 3’-junction of RPA-coated gap substrates but the specific interaction between RFC and RPA is dispensable. Rather, the ssDNA within gaps larger than 10 nt is a major factor that drives loading and stabilization of PCNA-RFC complexes. At an RPA-coated gap, stable PCNA-RFC complexes are on path for recruitment of Pol δ to PCNA, and these complexes are active for gap-filling DNA synthesis. Finally, we show that the combined activities of PCNA, Pol δ synthesis and Pif1 unwinding must be coordinated during strand displacement synthesis.

## Results

### PCNA is preferentially loaded at the 3’-junction of an RPA-coated ssDNA gap

To visualize PCNA loaded at an individual RPA-coated ssDNA, we performed single molecule experiments that combine optical trapping with confocal imaging. To that end, a doubly biotinylated ∼17.8 kb dsDNA molecule, with two nicks to generate a 5 knt central ssDNA gap, was tethered between two beads held in the optical traps (Supplementary Figure S1A). The substrate is labeled with an ATTO 647N fluorophore on the dsDNA at the 3’-edge of the ssDNA gap, allowing determination of the orientation of the tether (Figure 1A). After formation of the ssDNA gap, the fluorophore was purposefully photobleached before further use. The tether was then moved into a dedicated channel to allow coating of the ssDNA with MB543-labeled RPA (green) and imaged to estimate the length of the ssDNA gap (∼2.3 μm, Supplementary Figure S1B and S1C). Finally, the tether was moved to a channel where AATOM 647N-PCNA (red) was loaded by RFC and ATP (see Materials and Methods).

**Figure 1.**
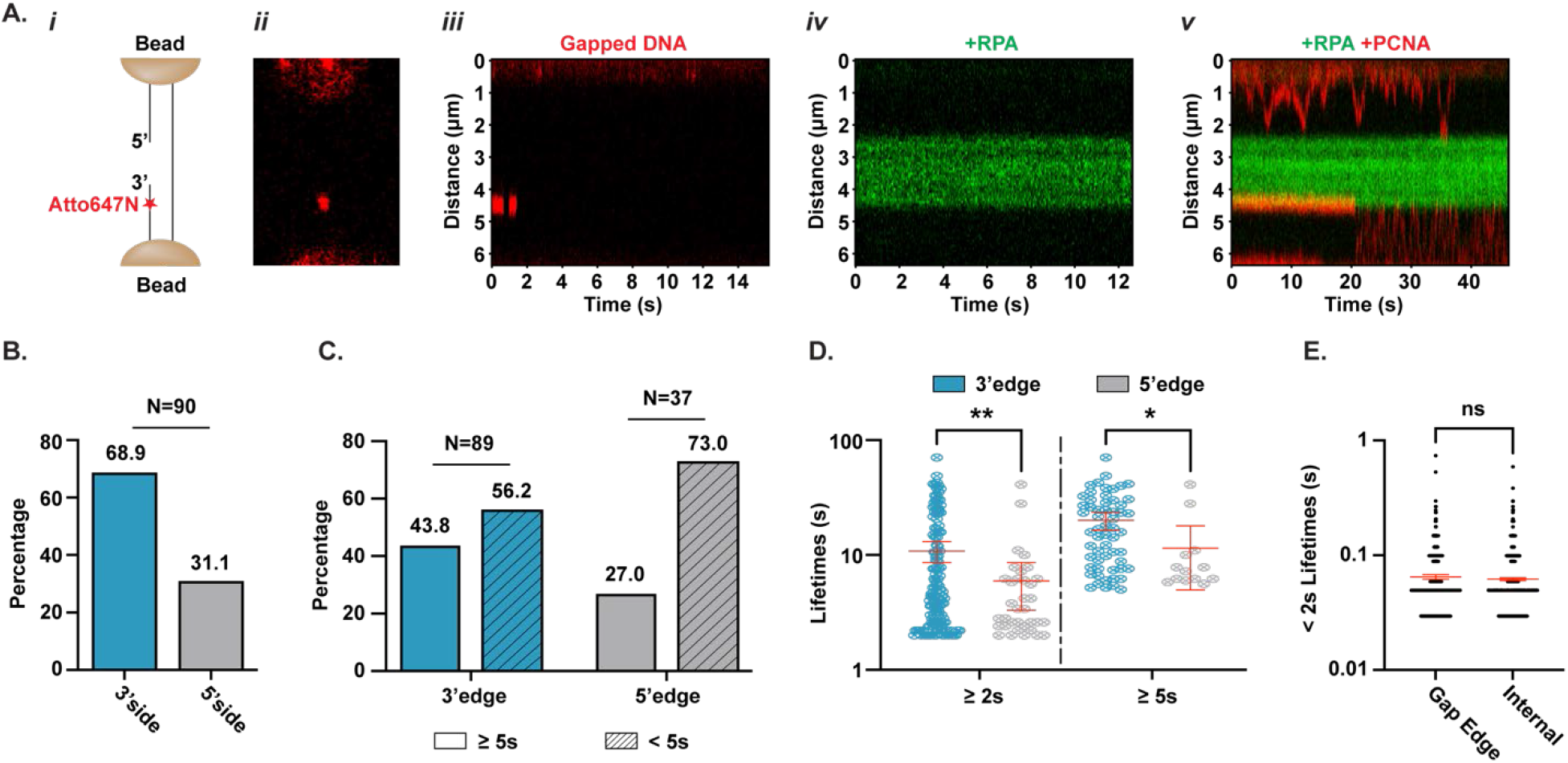
PCNA is loaded and stabilized preferentially on the 3’-edge of an RPA-coated gap. **A)** Experimental flow: i) a DNA with a centrally positioned 5 knt ssDNA gap, labeled with an ATTO 647N fluorophore on the 3’ side, was tethered between two optically trapped beads; ii) scanned image showing the edges of the two beads and the position of the fluorophore (red dot) along the tethered DNA; iii) kymograph showing photobleaching of the fluorophore; iv) kymograph of the MB543-RPA-coated gap (in green); v) kymograph displaying AATOM 647N-PCNA (in red) that was loaded by RFC at either side of the gap. **B)** Percentage of PCNA molecules loaded on the 3’ or 5’ side of the gap. **C)** Percentage of PCNA loading events at the 3’ or 5’ edge of the gap, classified either as long- (≥ 5s, solid) or short-lived (<5s, shaded) events based on a 5s cutoff. **D)** Lifetimes of long-lived events at the 3’ or 5’ edges, classified with either a 2s or 5s cutoff. **E)** Short-lived lifetimes (<2s cutoff) measured at the edges of the RPA-coated gap versus those measured at an internal position along the duplex DNA. Error bars in D) and E) represent the mean ± 95% confidence interval and statistical significance was determined using two-tailed unpaired Welch’s t test.

Counting of the RFC-dependent PCNA loading events detected at either the 5’- or 3’-side of the gap indicates a significant preference for PCNA loading on the dsDNA 3’ to the gap (Figure 1B). PCNA loaded at either side of the RPA-coated gap displays three classes of behavior: “diffusing”, and both “long”-, and “short”-lived states at the edges of the gap. PCNA diffuses along the dsDNA of the substrate with a diffusion coefficient of 0.62 ± 0.05 μm^2^/s (Supplementary Figure S3A and S3B), which is consistent with previous reports (15, 57). However, of interest here are the two classes of PCNA behavior at the edges of the RPA-coated gap. For the purpose of this paper, we take these edges as representing the 3’- and 5’ss/dsDNA junctions. On the one hand, analysis of the lifetimes of PCNA loaded at these junctions (Supplementary Figure S2) shows that the fraction of “long”-lived states is larger at the 3’-junction than at the 5’-junction, while the “short”-lived states dominate at the 5’-junction (Figure 1C). This observation is independent of how the lifetimes of these states were defined (Supplementary Figure S3C). Furthermore, the lifetime of the “long”-lived states at the 3’-junction is on average longer than that at the 5’-junction (Figure 1D). Taken together, these observations strongly suggest not only that PCNA is preferentially loaded at the 3’-junction of the gap, but also that the loaded PCNA is further stabilized at this end. On the other hand, the lifetimes of the “short”-lived states are the same at either end of the gap (Supplementary Figure S3D), suggesting that these do not represent stabilizing interactions with the ends of the gap. Indeed, these fast lifetimes are essentially the same as the ones calculated at an internal position along the dsDNA (Figure 1E), where PCNA is undergoing 1D diffusion. Thus, we conclude that the “short”-lived states at the edges reflect the diffusing mode of PCNA.

Next, we sought to define what drives the preferential stabilization of the “long”-lived state of PCNA at the 3’-edge of the RPA-coated gap. While PCNA and RPA do not interact directly, RFC4 has been shown to bind to RPA1 in *Saccharomyces cerevisiae* (58) and human RFC has been shown to bind human RPA (59). Moreover, yeast RFC binding to primer–template junctions is stimulated by yeast RPA binding to ssDNA, and this stimulation is dependent on RFC1 (8). Furthermore, human RPA specifically stimulates human RFC activity on primer-templates, when compared to yeast RPA (60). Thus, we examined the species-specific stimulation of yeast RFC-dependent PCNA loading on gaps coated with either labeled human RPA or *E. coli* SSB. Preferential loading of PCNA at the 3’-junction of the gap is maintained in either case (Figure 2A). This observation strongly suggests that while the RFC-RPA interaction may be important for localization of RFC at an RPA-coated gap, this interaction is not sufficient to drive the preferential loading and stabilization of loaded PCNA at the 3’-edge of the gap.

**Figure 2.**
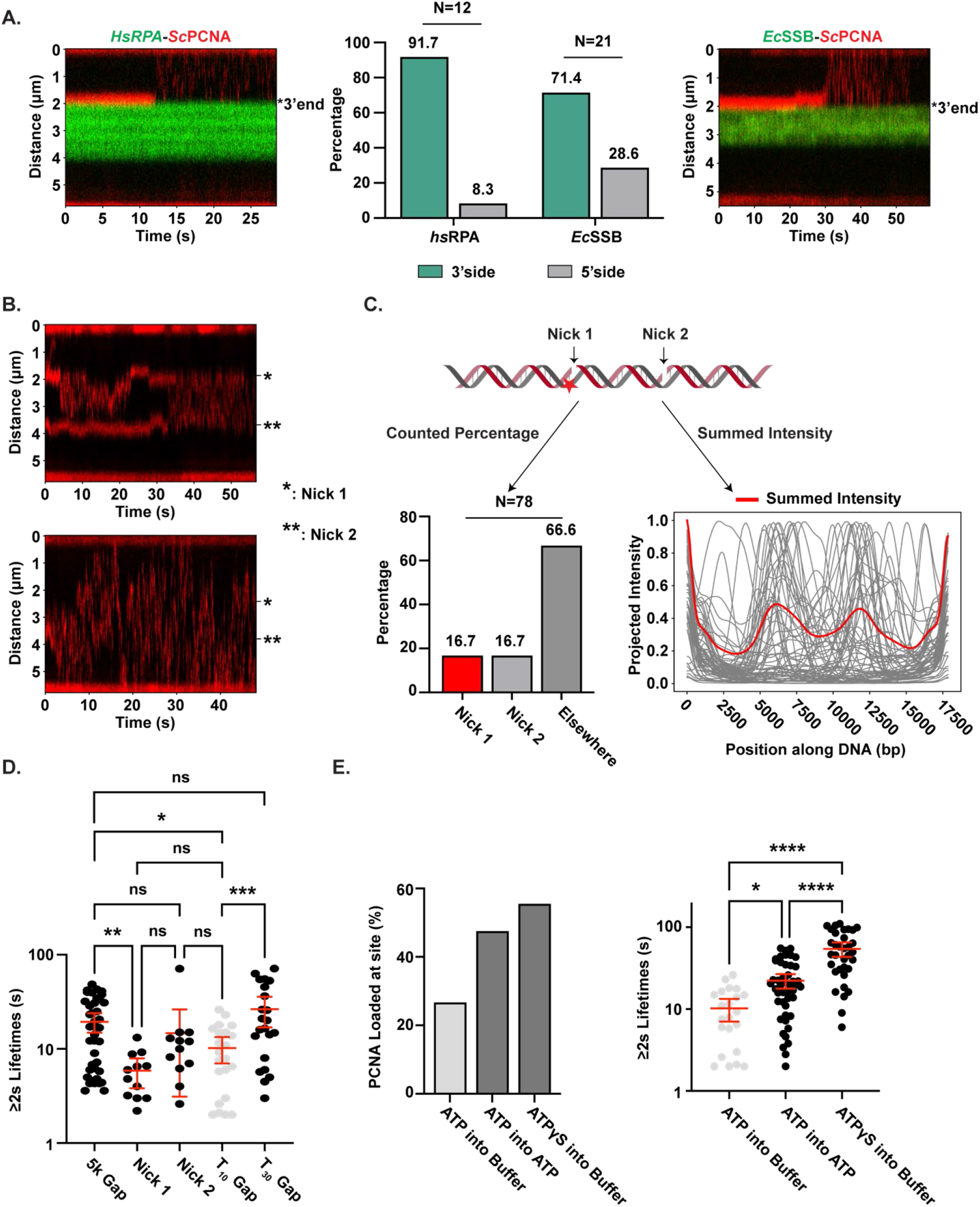
PCNA loading and stability is driven by RFC interaction with the ssDNA of the junction. **A)** *Sc*PCNA is loaded on *Hs*RPA- or *Ec*SSB-coated gapped substrates (left and right kymographs, respectively). The bar chart in the middle shows the percentage of *Sc*PCNA loaded on either side of *Hs*RPA- or *Ec*SSB-coated gapped gap. **B)** Representative kymographs of PCNA loaded onto a doubly nicked substrate. **C)** Schematic of the substrate showing Nick 1 in close proximity to the ATTO 647N fluorophore (Top). The bottom left bar chart shows the percentage of PCNA loaded at Nick1, Nick2 or somewhere else along the substrate. The bottom right graph shows the normalized intensity projected along the time axis of 61 kymographs of loaded PCNA (grey) plotted against the position along DNA. The normalized summed intensity of the kymographs is overlaid in red. **D)** Lifetimes of long-lived loaded PCNA at the 3’-edge of the RPA-coated 5k gap, Nick1 or Nick2 of nicked substrate, and at a T_10_ or T_30_ gap. **E)** The left panel shows the percentage of PCNA loaded at a T_10_ gap by RFC in the presence of either 1 mM ATP or 0.5 mM ATPγS, after the tethered DNA was moved into a channel with buffer only or 1 mM ATP. The corresponding lifetimes of the long-lived loaded PCNA are shown in the left panel. Error bars in D) and E) represent the mean ± 95% confidence interval and statistical significance was determined using ordinary one-way ANOVA.

### Interaction of RFC with ssDNA of the gap drives loading and stability of PCNA

If not via protein-protein interactions, the observed loading and stabilization of PCNA at the 3’-edge of the RPA-coated gap must be driven by one or more features of the 3’ ss/dsDNA junction. Consistent with its involvement in base excision repair (61), which is initiated at nicked DNA at sites of base damage, PCNA is known to also load at nicks (62, 63), which we have recapitulated in this study. First, we eliminated the possibility that the ATTO 647N fluorophore, positioned ∼ 170 nt from the 3’-junction affected PCNA loading or stabilization, as evidenced by the equal contributions of both nicks (Figure 2B-D, further explanation in Supplementary Information). We note that enrichment of loaded PCNA observed at the nicks relative to any other position along the DNA is higher than what would be expected for random binding (∼ 16% in Figure 2C *vs*. 3% in Supplementary Figure S4, respectively), suggesting that the nick does impart some level of preference for PCNA loading. However, once transiently loaded at the nick, PCNA undergoes fast 1D diffusion along the entire substrate (Supplementary Figure S3B). RFC itself binds with different affinities to nicks and small gaps, independent of RPA (54), and we take this as a first indication that the PCNA enrichment we observe at the nick relative to any other position along the DNA represents RFC-PCNA complexes.

Because of the reported higher affinity of RFC for small gaps compared to nicks (54), we tested whether small gaps would further increase the loading preference of PCNA or its stability. For this, we performed RFC-dependent PCNA loading on DNA tethers that contain a small 10 nt gap (T_10_) in the middle of the substrate. The frequency of PCNA loaded at the center of the tether is higher than expected for random loading (Supplementary Figure S6C), strongly suggesting a preference for loading at the T_10_ gap relative to any other position along the ∼12.6 kbp tether. However, the frequency and lifetime of PCNA loaded at this site are similar to the ones observed at nicks (Figure 2D). Thus, our data suggest that a T_10_ gap is too short to load an RFC-PCNA complex that is significantly more stable than the one loaded at a nick. Indeed, the lifetime of RFC-PCNA loaded at the T_10_ gap is shorter than the one observed at the RPA-coated gap (Figure 2D). The situation is different when the gap is extended from 10 to 30 nts. The frequency of PCNA loading at this larger gap is slightly higher and, importantly, the lifetime of RFC-PCNA at the T_30_ gap is longer than at the T_10_ gap, indicating further stabilization of the loaded complex by the additional length of ssDNA (Figure 2D and Supplementary Figure S6C). Furthermore, the lifetime of RFC-PCNA at the T_30_ gap is similar to the one at the 3’-edge of the RPA-coated gap (Figure 2D). Thus, the data suggest that at the RPA-coated gap, the RFC-PCNA complex is possibly stabilized by interaction with available free ssDNA at the ss/dsDNA junction rather than by specific interactions with RPA.

We note that in the former experiments the RFC- and ATP-dependent loading of PCNA was performed in one channel of the flow chamber and then the tethered DNA was moved into a separate observation channel, devoid of both RFC and ATP. Thus, the stability of the loaded RFC-PCNA complex is a combination of the lifetime of RFC bound to DNA and the rate of ATP dissociation from RFC and/or hydrolysis. Dissociation of RFC due to ATP dissociation and/or hydrolysis would result in an increase in the probability for PCNA to enter its 1D diffusion mode, thereby decreasing the fraction of stably loaded complexes and the apparent preference for the 3’-edge. To test this, we performed experiments in which after RFC-dependent PCNA loading, the tethered DNA was moved into an observation channel that contained excess ATP, instead of buffer only. In this experimental configuration, we observed an increased fraction of longer-lived PCNA loaded at the T_10_ gap (Figure 2E), consistent with excess ATP stabilizing the RFC-PCNA complex by suppressing RFC unloading due to ATP dissociation and/or hydrolysis. This conclusion is further corroborated by RFC-dependent PCNA loading performed in the presence of ATPγS, a slowly hydrolysable ATP analog that supports PCNA loading but greatly slows RFC release from the complex (8, 64). When ATP hydrolysis is suppressed, a high percentage of stably loaded PCNA is observed at the T_10_ gap (Figure 2E). Because excess ATP or ATPγS suppress RFC dissociation from DNA caused by ATP dissociation and/or hydrolysis, these observations reinforce the conclusion that PCNA that is stably loaded at a nick or gap must still be in complex with RFC, while PCNA undergoing 1D diffusion is not. This observation further supports the conclusion that RFC interaction with the ssDNA at the ss/dsDNA junction is a major driver of the stability of the loaded RFC-PCNA complexes.

### Pol δ stabilizes PCNA loaded at the 3’-junction to form a complex primed for DNA synthesis

During DNA replication, Pol δ binds to PCNA loaded at a 3’-junction to form a complex that carries out gap DNA synthesis while displacing bound RPA (28). In the process, RFC is expected to leave its complex with PCNA to allow for Pol δ binding (7). These early studies were carried out with a form of RFC in which the N-terminal BRCT domain of the Rfc1 subunit has been removed (∼550 aa in human, ∼270 aa in yeast RFC), because the Rfc1-NTD showed non-specific DNA binding and a moderate inhibition of DNA synthesis by Pol δ (7, 56, 65). We are also using the truncated form of RFC in this paper. However, recent work using full-length RFC has argued for an alternative model in which RFC-PCNA-Pol δ can form a ternary complex that carries out processive DNA synthesis (12).

Notwithstanding, if RFC unloads from PCNA, one question is whether the simultaneous binding of PCNA and Pol δ to the 3’-junction is sufficient to stabilize PCNA at that end, before the onset of DNA synthesis (e.g. in the absence of dNTPs). To answer this question, we performed RFC-dependent loading of PCNA at an RPA-coated gap in the presence of Pol δ, followed by observation of the trapped DNA tether in a channel devoid of any additional RFC, PCNA or Pol δ· Under these conditions, the preference of PCNA loading at the 3’-side of the RPA-coated gap is maintained (Figure 3A); in addition, Pol δ does not appear to alter significantly the balance of the overall loading events at either side of the gap. Rather, Pol δ affects the distribution of “long”- and “short”-lived states observed at either edge. While at the 5’-edge Pol δ does not alter the fraction of “long”- to “short”-lived states, at the 3’-edge the fraction of “long”-lived states increases in the presence of Pol δ, independent of the time cutoff used (Figure 3B).

**Figure 3.**
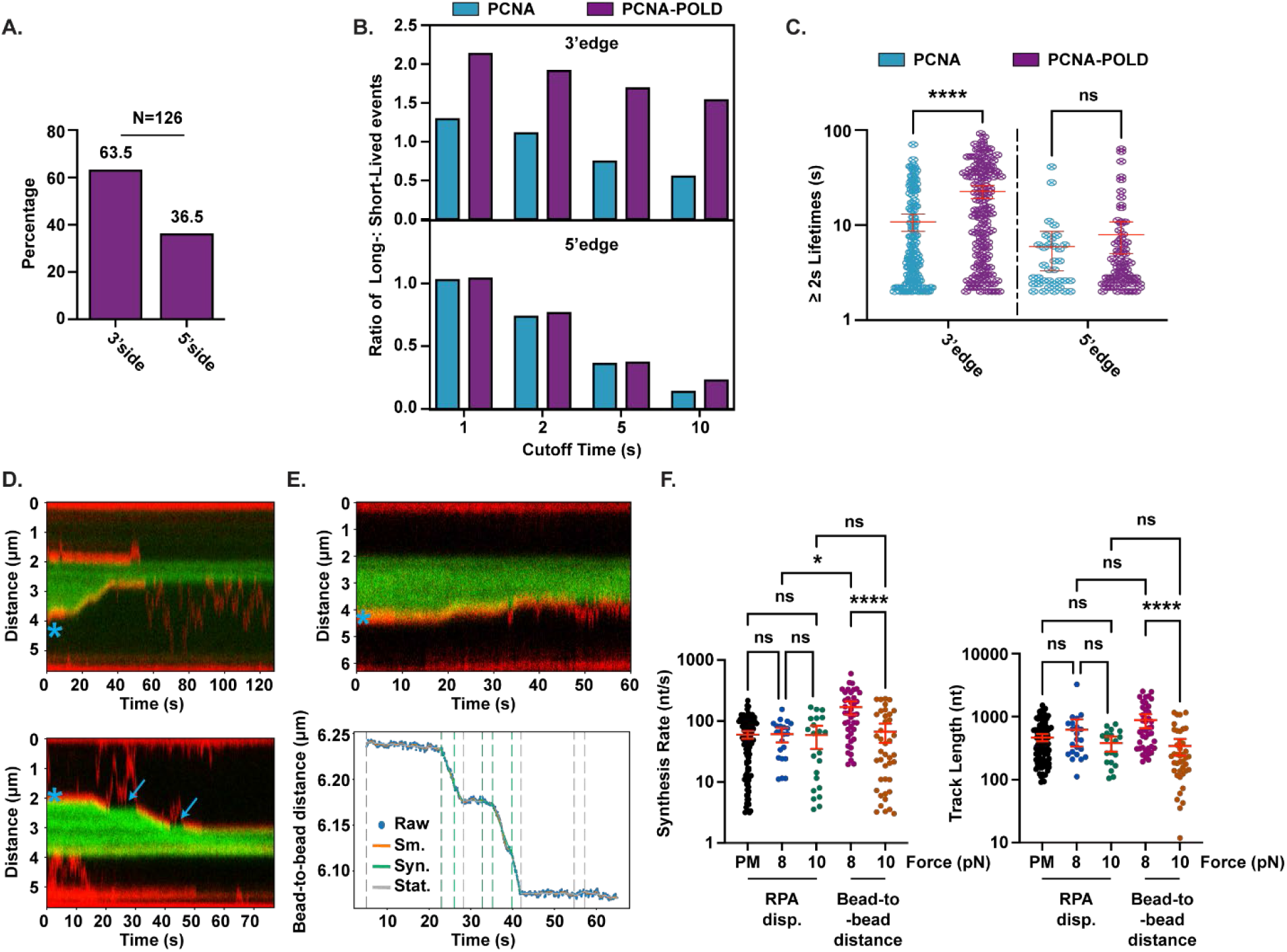
Loaded PCNA is stabilized by Pol δ and the complex catalyzes gap-filling synthesis of the RPA-coated gap. **A)** Percentage of PCNA loaded in the presence of Pol δ on either side of the RPA-coated gap. **B)** Ratio of the percentage of long-lived to short-lived PCNA loading events at the 3’- (top) or 5’-edge (bottom) of the gap, calculated with different cutoff times in the presence (purple) or absence (blue) of Pol δ. **C)** Lifetimes of long-lived loaded PCNA events at the 3’- or 5’-edge, in the presence (purple) or absence (blue) of Pol δ. Statistical significance was determined using ordinary one-way ANOVA. **D)** Representative kymographs showing gap-filling synthesis by Pol δ (in passive mode), as monitored by displacement of RPA (green) and concomitant directional PCNA movement along the 3’-edge of the gap (marked by an asterisk*). The arrow marks an example of pausing in RPA displacement accompanied by PCNA diffusion away from the edge. **E)** Top panel: a kymograph of RPA displacement and PCNA directional movement under force-clamp (10 pN). Bottom panel: corresponding bead-to-bead distance versus time curve. The decrease in distance is due to more ssDNA being converted to dsDNA as synthesis proceeds. The raw data (blue) were smoothed (Sm.) and then segmented (see Materials and Methods). **F)** Gap-filling synthesis rate (Left) and track length (Right) calculated either by RPA displacement or bead-to-bead distance changes (PM: passive mode) (see Supplementary Fig. 5B, C and Materials and Methods). Statistical significance was determined using a Kruskal-Wallis test. Error bars in C) and F) represent the mean ± 95% confidence interval.

Similarly, the lifetime of the “long”-lived state in the presence of Pol δ is increased at the 3’- junction but not the 5’-junction (Figure 3C). Thus, we conclude that Pol δ leads to preferential stabilization of PCNA loaded at the 3’-junction, even without DNA synthesis. Interestingly, Pol δ also increases the lifetime of the “short”-lived states, both at the 3’- and 5’-edges (Supplementary Figure S3D and S3E). As already noted, these “short”-lived PCNA states at the edge of the RPA-coated gap represent the 1D diffusion mode of PCNA, rather than specific interactions with the junctions. Yet, Pol δ increases their lifetime. The increase of these “short” lifetimes may be rationalized by an overall decrease of PCNA 1D diffusion coefficient in the presence of Pol δ due to the increased size of the complex (Supplementary Figure S3B).

Importantly, independent of how the PCNA-Pol δ complex is formed at the 3’-junction, this is the complex that is active for DNA synthesis. No DNA synthesis into the RPA-coated gap can be detected in the absence of PCNA (Supplementary Figure S5E). However, in the presence of PCNA and dNTPs, DNA synthesis from the 3’-junction can be readily detected either as the directional displacement of the “green” labeled RPA from the gap and concomitant movement of the “red” labeled PCNA (in passive mode, Figure 3D and Supplementary Figure S5A) or as a change in RPA and PCNA signals with concomitant change in bead-to-bead distance due to the increased content of newly synthesized DNA (in force clamp mode, Figure 3E). Analysis using the change of the RPA signal as a proxy for DNA synthesis (Supplementary Figure S5B and S5C), indicates gap filling proceeds at an average rate of 64.3 ± 4.3 nt/s and with an average track length of 465.5 ± 31.5 nt (Figure 3F). These values do not change when the tethered DNA is held at constant force of 8 or 10 pN (Figure 3F), indicating that neither the rate of RPA displacement nor the amount that is being displaced are significantly force-dependent. That is not to say that DNA synthesis itself is not force-dependent. Indeed, both the rate and track length of DNA synthesis measured from bead-to-bead distance are significantly higher when the tethered DNA is held at a lower constant force (Figure 3F).

### Pol δ and Pif1 stabilize PCNA at the 5’-flapped substrate

During BIR and strand displacement DNA synthesis in general, loading of PCNA at a 3’- junction occurs in the context of a 5’-tail on the downstream duplex to be unwound, rather than an RPA-coated gap. Thus, we next tested the effect of a 5’-flap in the duplex downstream of a short T_10_ gap (T_10_ gap – T_25_ flap) on the RFC-dependent PCNA loading (Figure 4A). Surprisingly, the fraction of loaded PCNA events at this site is higher than at a simple T_10_ gap (Supplementary Figure S6C). An increase in the fraction of loaded PCNA is observed also when the gap is shortened to a T_1_ gap, pointing to a direct effect of the 5’-tail on loading efficiency (Supplementary Figure S6C). The length of the ssDNA gap still has a stabilizing role: the T_10_ gap – T_25_ flap increases the lifetime of PCNA more than the shorter T_1_ gap at the T_1_ gap – T_25_ flap site (Figure 4B). This reinforces the conclusion that RFC binding to the gap contributes to the stability of the RFC-PCNA complex. Importantly, the effect of the 5’-flap cannot be simply explained by the flap blocking PCNA diffusion past it. If this were the case, we would not expect an increase in either the frequency of loading or the lifetime of PCNA at this site. Rather, the stabilizing effect of the 5’- flap must be via direct interaction with either PCNA or RFC. We favor the latter and interpret these results as evidence that in addition to binding to the ssDNA in the gap, one or more subunits of RFC can engage in interactions with the flap.

**Figure 4.**
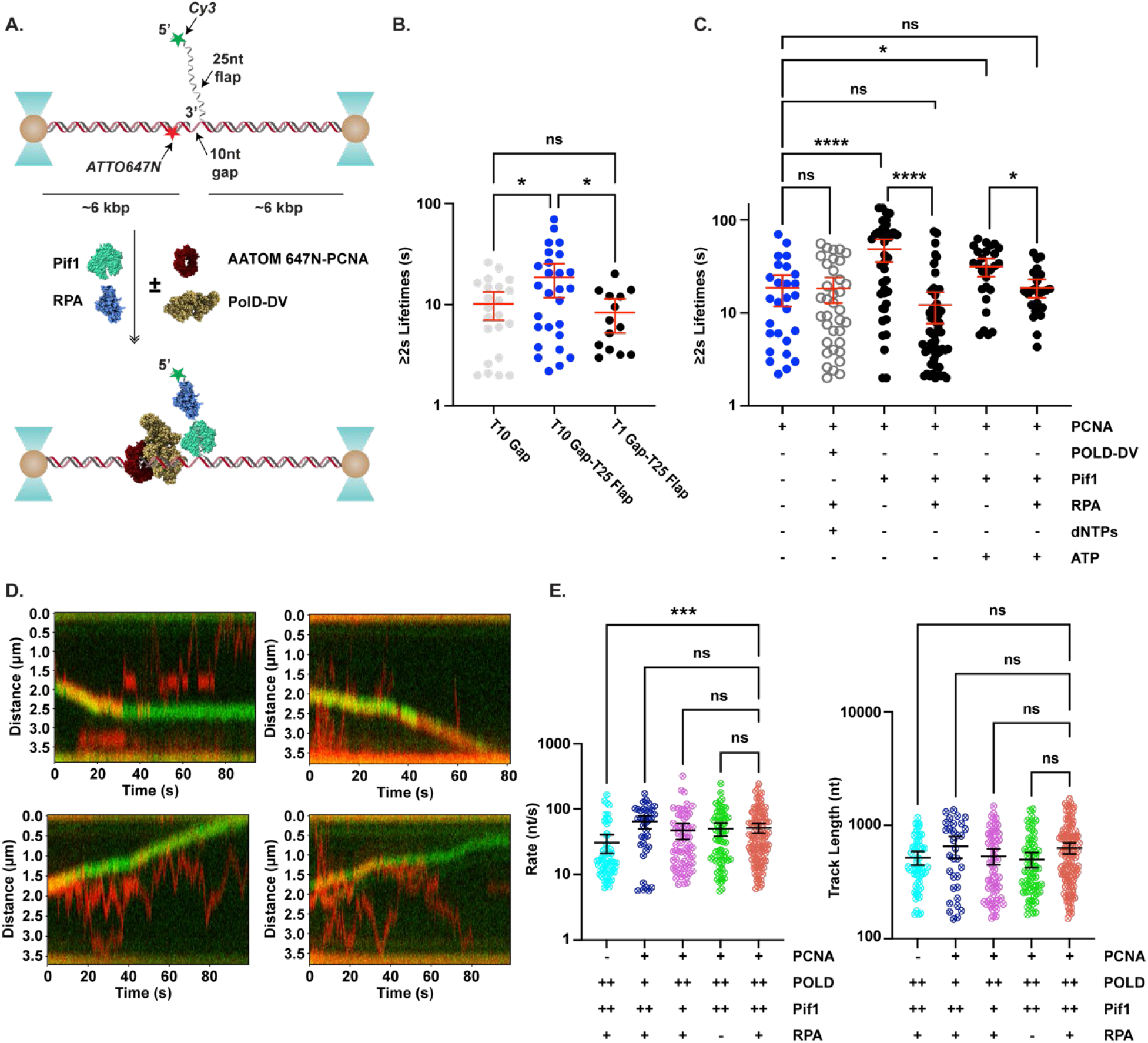
Pif1 is required for strand displacement DNA synthesis. **A)** Schematic of the experimental flow for replication experiments using a DNA tether that contains a centrally located gap and a 5’ Cy3- labeled flap. **B)** Lifetimes of long-lived loaded PCNA events at T_10_ gap, T_10_ gap with a T_25_ 5’-flap or T_1_gap with T_25_5’-flap. **C)** Lifetimes of long-lived loaded PCNA events in the presence of different combinations of Pol δ, Pif1, unlabeled RPA, dNTPs and ATP. **D)** Representative kymographs showing strand displacement DNA synthesis as monitored by the directional movement of the Cy3-labeled 5’flap (green) due to the increase of displaced ssDNA and the movement of AATOM 647N-labeled PCNA (red) along the green signal or diffusing away. **E)** Strand displacement synthesis rate (Left) and track length (Right) in the presence of different combinations of proteins. Statistical significance in B) and C) was determined using ordinary one-way ANOVA and Kruskal-Wallis test in E). Error bars in B), C) and E) represent the mean ± 95% confidence interval.

Next, we tested whether the 5’-flap alters the PCNA-stabilizing role of Pol δ that we observed at the RPA-coated gap. In the context of a model in which RFC unloads from PCNA before Pol δ binding, Pol δ interaction with the 3’-junction and/or template must substitute for RFC in stabilizing PCNA at the site, preventing PCNA from sliding away from the 3’-junction and entering its 1D diffusing mode. In support of this model, Pol δ primed for DNA synthesis (+dNTP) does not increase the fraction of 1D diffusing PCNA molecules (Figure 4C and Supplementary Figure S6D).

PCNA-Pol δ complexes do not catalyze efficient strand displacement DNA synthesis, slowing down >100 fold from the elongation mode on a ssDNA template (28). During BIR the activity of the accessory helicase Pif1 is required to perform efficient strand displacement synthesis (36, 37, 45). Pif1 interacts with PCNA (36, 37, 43), yet this interaction appears to be dispensable for its in vitro activities (40, 43) but required for efficient replication in cells (36, 37, 43). We note that Pif1 slows down the 1D diffusion coefficient of PCNA, which is an indication of their interaction on the DNA (Supplementary Figure S3B). Next, we tested whether the addition of Pif1 alters the stability of PCNA loaded at the 3’-junction of the T_10_ gap – T_25_ flap structure. In the presence of Pif1, the fraction of PCNA loaded at the site is slightly higher (Supplementary Figure S6D). More importantly, Pif1 increases significantly the stability of loaded PCNA (Figure 4C). When RPA, which can compete with Pif1 for binding the 5’-flap, is also present, the lifetime of PCNA on DNA is significantly lowered (Figure 4C), while the fraction of PCNA loaded at the site is little affected (Supplementary Figure S6D). These observations suggest that the stabilizing effect of Pif1 on PCNA is mediated by the ability of Pif1 to bind to the 5’-flap. The same is the case in the presence of ATP, indicating that the Pif1-PCNA stabilizing effect occurs also with Pif1 primed for unwinding (Figure 4C and Supplementary Figure S6D).

### Pif1 is required for strand displacement DNA synthesis

Stabilization of PCNA at the 3’-junction by interaction with Pif1 bound to the 5’-flap of the substrate suggests a direct coupling between PCNA, Pif1 and the DNA substrate to be unwound. Even so, no unwinding of the substrate by Pif1 is detected when monitoring the position of the Cy3-labeled 5’-flap (Figure 4A). An increase in the length of the 5’-flap due to unwinding is expected to compact the ssDNA because of its low persistent length, and result in directional movement of the green signal along the tethered DNA. This is not observed (Supplementary Figure S6A). Also, no strand displacement DNA synthesis by PCNA-Pol δ is detected (Supplementary Figure S6B).

The situation is different when, after PCNA loading, the DNA tethers were moved into a channel that contains Pol δ, Pif1, RPA, dNTPs and ATP. Under these conditions, unidirectional movement of the signal from the labeled 5’-flap and labeled PCNA can be observed (Figure 4D). In the presence of PCNA, both the rate and track length of strand displacement DNA synthesis are little affected by a 10-fold change in the concentration of either the helicase or the polymerase, indicating that neither is limiting under these conditions (Figure 4E). The rate of strand displacement DNA synthesis (51.8 ± 4.4 nt/s) is only slightly slower than the rate of gap-filling synthesis, suggesting that the rate limiting step may be dominated by the rate of DNA synthesis by the polymerase rather than opening of the downstream duplex by the helicase. Interestingly, the average track length of strand displacement synthesis is 632.6 ± 35.8 nts, higher than that of gap-filling synthesis (465.5 ± 31.5 nt). Furthermore, the presence of RPA does not alter the rate and the track length (Figure 4E), indicating that albeit it can compete with Pif1 for binding to the 5’-flap, this is not sufficient to block access by Pif1. Surprisingly, PCNA is not strictly required for strand displacement DNA synthesis (Figure 4E and Supplementary Figure S6E), suggesting that its interaction with neither Pol δ nor Pif1 is necessary. In the absence of PCNA, the rate of strand displacement DNA synthesis is slower by about two-fold, but neither the track length nor the pause time between synthesis events are affected (Figure 4E and Supplementary Figure S6F).

## DISCUSSION

RFC-dependent loading of PCNA at 3’-junctions to be extended by Pol δ is a crucial step in lagging strand DNA replication and in DNA repair via BIR. Biochemical studies have shown that RFC preferentially loads PCNA at a 3’ ss/dsDNA junction (5, 8, 62). However, several questions remain. For example, what specific features of the junction make it a preferential loading site? Or, once the site is recognized, what determines the lifetime of PCNA at the site relative to any other position along the DNA? Here, we used single molecule confocal imaging of tethered DNA to visualize loaded PCNA, disentangle different modes of interaction, and dissect the determinants for the preferential RFC-dependent PCNA loading and stability.

We showed that 1D diffusion of PCNA is a key dynamic mode of interaction with DNA, as previously demonstrated (15). In a “PCNA-centric” view of the PCNA loading process, this mode of interaction may play a role in the ability of PCNA to form a stable complex with Pol δ at the 3’- junction. Diffusing PCNA may be able to interact with a Pol δ molecule already bound to the 3’- junction, and this interaction may result in stabilization of a PCNA-Pol δ complex primed for DNA synthesis. In this pathway, stable complexes would result from transient interaction with Pol δ pre-bound at the site, rather than PCNA being at the site first and serving as a loading platform for Pol δ. Alternatively, it is tempting to speculate that binding of Pol δ to PCNA while it is diffusing along the dsDNA, could be a means to “transport” the polymerase to the 3’-junction. In this regard, we note that the 1D diffusion coefficient of PCNA is reduced in the presence of Pol δ, and this slowing down of the complex may provide an alternative means to find and interact with the 3’- junction. The decrease in the 1D diffusion coefficient is consistent with PCNA forming complexes with the large Pol δ (66) that diffuse slower on DNA, as observed also upon interaction with Pif1 (Supplementary Figure S3B).

Alternatively, we favor a more “RFC-centric” view, where we propose that the 1D diffusing mode of PCNA is a competing pathway to the stable and productive loading of PCNA at the 3’- junction driven by RFC. We showed that PCNA can be loaded in a stable mode at an available 3’-junction, with “long”-lived states that we ascribe to PCNA being still in complex with RFC. This conclusion is supported by experiments performed either in excess ATP, to slow down ATP dissociation/hydrolysis from RFC, or ATPγS, a slowly hydrolysable ATP-analog, that supports loading but slows down RFC release. In either condition the lifetime of the PCNA at the 3’-edge of the gap increases. Because only RFC binds ATP, we conclude that the “long”-lived mode of PCNA at the 3’-junction represents stably bound RFC-PCNA complexes. Further support for this conclusion comes from the fact that RFC and PCNA have been visualized simultaneously at the edge of an RPA gap (12). PCNA can also exist in “short”-lived states at the edge of an RPA-coated gap; however, we ascribe these to PCNA rings that have been released from RFC and are now freely diffusing along dsDNA. Importantly, we showed that Pol δ preferentially stabilizes PCNA at the 3’-edge of an RPA-coated gap, without an increase in the fraction of 1D diffusion mode of PCNA. We interpret this as a result of Pol δ being able to stabilize PCNA at the 3’-junction by replacing the stabilizing interactions provided by RFC, consistent with the idea of a hand-off. In support of this conclusion, we note that ternary complexes of RFC-PCNA-Pol δ have only been observed for the full-length RFC1, and not with its N-terminus truncated variant (12), which was used in our studies.

Our data also indicate that a simple nick within a large dsDNA is sufficient to provide a preferred RFC-PCNA loading site, albeit the loaded complexes are not as stable as the ones formed at gaps. Biochemical and structural studies have shown that small gaps are preferred RFC interaction sites (54). Here, we showed that while there is some preference for a small 10 nt gap, it is only when the gap is extended to 30 nts that the stability of the loaded RFC-PCNA complexes increases significantly. Indeed, these complexes are as stable as the ones formed at an RPA-coated gap. Similar stability at these two different sites is consistent with our findings that loading of PCNA at RPA-coated gaps does not seem to be strictly driven by specific protein-protein interaction. Preferential loading at the 3’-edge is maintained also when the gap is coated with either *hs*RPA or *Ec*SSB. Thus, we conclude that a ssDNA gap longer than 10 nts is a major factor in stabilizing loaded RFC-PCNA complexes. A downstream 5’-ssDNA flap in the context of a T_10_gap is also sufficient to stabilize the RFC-PCNA complexes, suggesting that more than one RFC binding sites may interact with different ssDNA regions at the site. However, even in the presence of the 5’-flap, if the gap size is shortened to a T_1_, the stabilizing effect is lost. Thus, the ssDNA in the gap remains a major driver of stability.

The “long”-lived states of PCNA-Pol δ formed at the 3’-edge of the RPA-coated gap are primed for DNA synthesis. Stretches of DNA synthesis are interspersed with pauses of no activity, whose duration is independent of whether RPA displacement or the bead-to-bead distance is monitored (Supplementary Figure S5F). Therefore, we parsed out these pauses and analyzed only regions of active DNA synthesis or RPA displacement. For the latter, the measured rates and track lengths appear to be independent of force. However, when DNA synthesis is measured from the bead-to-bead distance both the rate and the track length appear to decrease as the applied force is increased. This suggests that, similarly to other polymerases (67-69), the DNA synthesis activity of Pol δ is force sensitive. Interestingly, at the lower force of 8 pN the rate of DNA synthesis measured by the change in bead-to-bead distance is higher than when measured by imaging RPA displacement (Figure 3F). This may be due to the two signals reporting on two different phenomena, with RPA displacement including the additional steps of protein removal that may be partially limiting to DNA synthesis.

We note that several of the pauses in the signal for RPA displacement coincide with PCNA diffusing away from the edge of the RPA-coated gap (arrows in Figure 3D and Supplementary Figure S5A). Because Pol δ does not carry out evident gap-filling synthesis in the absence of PCNA (Supplementary Figure S5E), loss of PCNA-Pol δ interaction and diffusion away of PCNA from Pol δ at the 3’-junction would be sufficient to result in a pause of DNA synthesis. Alternatively, the observed pausing could simply be due to Pol δ dissociation from the 3’-junction, followed by PCNA entering its 1D diffusion. In this latter case, in the presence of free Pol δ in the observation channel, a change in Pol δ concentration should result in a change in the average pause time. However, for strand displacement DNA synthesis a 10-fold change in enzyme concentration does not affect the pause time between synthesis events either (Supplementary Figure S6F). Thus, we interpret the observed pauses during gap filling as originating from the polymerase either stalling or losing its interaction with PCNA.

In cells, the activity of Pif1 and its interaction with PCNA are important for efficient BIR (36, 37, 45). PCNA loaded at the 3’-junction and the 5’-flap downstream of the 10 nt gap may provide binding sites for the Pif1 helicase and allow processive DNA unwinding of the duplex to be replicated during strand displacement DNA synthesis. In the presence of Pif1, we found that loaded PCNA is further stabilized at the 3’-junction of the 10 nt gap with a 5’-flap, an indication that PCNA-Pif1 may form a complex on DNA. However, formation of a PCNA-Pif1 complex at the junction is still not sufficient to elicit significant amount of DNA unwinding. Rather, DNA synthesis on the opposite strand is required. This is consistent with our findings that the activity of Pif1 helicases is dominated by a repetitive cycle of DNA opening and closing, leading to limited net DNA unwinding (41). As such, unwinding is limited to the transient opening of short stretches of DNA of ∼ 20 bp (41, 42). These events are below spatial resolution when monitoring the 5’-flap position along the DNA tether. Nonetheless, we expect these short DNA opening events to be sufficient for the polymerase to copy the template that is transiently made available by the helicase. Indeed, we find that the rate of strand displacement DNA synthesis is limited by the rate of DNA synthesis of Pol δ and not unwinding by Pif1.

The combination of Pol δ and Pif1 activities is needed to observe strand displacement DNA synthesis; however, PCNA does not appear to be strictly needed in the process. Despite Pif1 leading to stabilization of loaded PCNA, this interaction is not required for the activity of the polymerase. While PCNA-Pif1 interaction is important for activity in cells (37), its requirement can be partially bypassed in vitro (41). This observation is consistent with our proposal that the PCNA- Pif1 interaction may be used to localize Pif1 at the site of action, rather than to modulate its enzymatic activities (43). However, the situation is different for Pol δ. In the presence of PCNA, the rate of strand displacement DNA synthesis is faster. Stimulation by PCNA of the DNA synthesis rate of Pol δ is consistent with our previous findings that binding of Pol δ to PCNA increases its catalytic rate (70).

Finally, the dispensability of PCNA-Pol δ interaction during strand displacement DNA synthesis (Figure 4E) is very different from what is observed in gap-filling DNA synthesis, where PCNA is required (Supplementary Figure S5E). We speculate that during gap-filling synthesis in the absence of PCNA, the encounter of Pol δ with RPA would stall the polymerase and lead to its dissociation. PCNA interaction would be needed to stabilize Pol δ on DNA, making PCNA required for efficient displacement of bound RPA. However, during strand displacement DNA synthesis Pif1 would open the DNA in front of Pol δ, and the polymerase would be free to replicate a “naked” template, thereby bypassing the requirement for its interaction with PCNA. This scenario would also explain the observation that the processivity during strand displacement DNA synthesis appears to be higher than during gap-filling. Even in the presence of PCNA, encounter with an RPA to be displaced during gap-filling may still result in dis-engagement of Pol δ from the 3’- junction while still maintaining interaction with PCNA, thus reducing processivity. This may not be the case during strand displacement DNA synthesis, where Pol δ would not need to displace RPA as Pif1 opens the downstream duplex.

## Materials and Methods

Full description of Materials and Methods is included in Supplementary Information.

Briefly, DNA substrates used in this work are Lumicks Biotinylated DNA Hybrid and Lumicks DNA tethering kit. To generate the different DNA structures, the handles from the DNA tethering kit were ligated to short DNA substrates. For those, oligonucleotides were purchased from IDT and are listed in Table S1. Protein purifications followed previously published procedures. PCNA purification and labeling is detailed in Supplementary Information.

Single molecule experiments employed a Lumicks C-Trap setup with optical traps and confocal imaging. A flow cell with 5 barrierless channels housed the experimental procedure. Channels 1-3 were used to generate a DNA tether that was moved to channels 4 and 5 to bind different proteins. DNA tethers in the presence or absence of proteins were imaged with red and/or green lasers to construct scanned images and kymographs. These kymographs were analyzed using Python scripts to estimate gap length, lifetimes and diffusion coefficients. In addition, gap-filling synthesis and strand displacement synthesis were analyzed using a multi-segment fitting approach to identify stationary segments from those representing active synthesis.

## Acknowledgements

We would like to thank Dr. Kacey N. Mersch for continued discussion and training on the Lumicks C-TRAP; Dr. Alexander G. Kozlov for sharing labeled *Ec*SSB, and Dr. Timoth M. Lohman for discussing single molecule results. R.G. thanks Dr. Eric A. Galburt for discussions resulting in the right panel of Figure 2C.

## Supplementary Information

It contains Materials and Methods, and Supplementary Figures S1 to S6 and Table S1.

## References

1. Z. F. Pursell, I. Isoz, E. B. Lundström, E. Johansson, T. A. Kunkel, Yeast DNA polymerase epsilon participates in leading-strand DNA replication. Science 317, 127–130 (2007).

2. T. A. Kunkel, P. M. Burgers, Dividing the workload at a eukaryotic replication fork. Trends Cell Biol 18, 521–527 (2008).

3. P. M. Burgers, Polymerase dynamics at the eukaryotic DNA replication fork. J Biol Chem 284, 4041–4045 (2009).

4. G. Prelich et al., Functional identity of proliferating cell nuclear antigen and a DNA polymerase-delta auxiliary protein. Nature 326, 517–520 (1987).

5. T. Tsurimoto, B. Stillman, Functions of replication factor C and proliferating-cell nuclear antigen: functional similarity of DNA polymerase accessory proteins from human cells and bacteriophage T4. Proc Natl Acad Sci U S A 87, 1023–1027 (1990).

6. T. S. Krishna, X. P. Kong, S. Gary, P. M. Burgers, J. Kuriyan, Crystal structure of the eukaryotic DNA polymerase processivity factor PCNA. Cell 79, 1233–1243 (1994).

7. V. N. Podust, N. Tiwari, S. Stephan, E. Fanning, Replication factor C disengages from proliferating cell nuclear antigen (PCNA) upon sliding clamp formation, and PCNA itself tethers DNA polymerase delta to DNA. J Biol Chem 273, 31992–31999 (1998).

8. X. V. Gomes, P. M. Burgers, ATP utilization by yeast replication factor C. I. ATP-mediated interaction with DNA and with proliferating cell nuclear antigen. J Biol Chem 276, 34768–34775 (2001).

9. G. D. Bowman, M. O’Donnell, J. Kuriyan, Structural analysis of a eukaryotic sliding DNA clamp-clamp loader complex. Nature 429, 724–730 (2004).

10. C. Indiani, M. O’Donnell, The replication clamp-loading machine at work in the three domains of life. Nat Rev Mol Cell Biol 7, 751–761 (2006).

11. A. M. van Oijen, J. J. Loparo, Single-molecule studies of the replisome. Annu Rev Biophys 39, 429–448 (2010).

12. G. N. L. Chua et al., A non-catalytic role for RFC in PCNA-mediated processive DNA synthesis. Cell 189, 1124–1134.e1114 (2026).

13. N. Y. Yao, R. E. Georgescu, J. Finkelstein, M. E. O’Donnell, Single-molecule analysis reveals that the lagging strand increases replisome processivity but slows replication fork progression. Proc Natl Acad Sci U S A 106, 13236–13241 (2009).

14. J. S. Lewis et al., Single-molecule visualization of Saccharomyces cerevisiae leading-strand synthesis reveals dynamic interaction between MTC and the replisome. Proc Natl Acad Sci U S A 114, 10630–10635 (2017).

15. A. B. Kochaniak et al., Proliferating cell nuclear antigen uses two distinct modes to move along DNA. J Biol Chem 284, 17700–17710 (2009).

16. G. Maga et al., Okazaki fragment processing: modulation of the strand displacement activity of DNA polymerase delta by the concerted action of replication protein A, proliferating cell nuclear antigen, and flap endonuclease-1. Proc Natl Acad Sci U S A 98, 14298–14303 (2001).

17. P. Garg, P. M. Burgers, DNA polymerases that propagate the eukaryotic DNA replication fork. Crit Rev Biochem Mol Biol 40, 115–128 (2005).

18. R. Ayyagari, X. V. Gomes, D. A. Gordenin, P. M. Burgers, Okazaki fragment maturation in yeast. I. Distribution of functions between FEN1 AND DNA2. J Biol Chem 278, 1618–1625 (2003).

19. P. Garg, C. M. Stith, N. Sabouri, E. Johansson, P. M. Burgers, Idling by DNA polymerase delta maintains a ligatable nick during lagging-strand DNA replication. Genes Dev 18, 2764–2773 (2004).

20. L. Balakrishnan, R. A. Bambara, Okazaki fragment metabolism. Cold Spring Harb Perspect Biol 5 (2013).

21. L. Zheng, B. Shen, Okazaki fragment maturation: nucleases take centre stage. J Mol Cell Biol 3, 23–30 (2011).

22. D. J. Bentley et al., DNA ligase I null mouse cells show normal DNA repair activity but altered DNA replication and reduced genome stability. J Cell Sci 115, 1551–1561 (2002).

23. M. Kucherlapati et al., Haploinsufficiency of Flap endonuclease (Fen1) leads to rapid tumor progression. Proc Natl Acad Sci U S A 99, 9924–9929 (2002).

24. E. Larsen, C. Gran, B. E. Saether, E. Seeberg, A. Klungland, Proliferation failure and gamma radiation sensitivity of Fen1 null mutant mice at the blastocyst stage. Mol Cell Biol 23, 5346–5353 (2003).

25. R. S. Murante, L. Rust, R. A. Bambara, Calf 5’ to 3’ exo/endonuclease must slide from a 5’ end of the substrate to perform structure-specific cleavage. J Biol Chem 270, 30377–30383 (1995).

26. S. H. Bae, K. H. Bae, J. A. Kim, Y. S. Seo, RPA governs endonuclease switching during processing of Okazaki fragments in eukaryotes. Nature 412, 456–461 (2001).

27. J. B. Boule, V. A. Zakian, Roles of Pif1-like helicases in the maintenance of genomic stability. Nucleic Acids Res 34, 4147–4153 (2006).

28. J. L. Stodola, P. M. Burgers, Resolving individual steps of Okazaki-fragment maturation at a millisecond timescale. Nat Struct Mol Biol 23, 402–408 (2016).

29. M. L. Rossi et al., Pif1 helicase directs eukaryotic Okazaki fragments toward the two-nuclease cleavage pathway for primer removal. J Biol Chem 283, 27483–27493 (2008).

30. J. E. Pike, P. M. Burgers, J. L. Campbell, R. A. Bambara, Pif1 helicase lengthens some Okazaki fragment flaps necessitating Dna2 nuclease/helicase action in the two-nuclease processing pathway. J Biol Chem 284, 25170–25180 (2009).

31. M. A. Sparks, P. M. Burgers, R. Galletto, Pif1, RPA, and FEN1 modulate the ability of DNA polymerase delta to overcome protein barriers during DNA synthesis. J Biol Chem 295, 15883–15891 (2020).

32. X. Li, C. M. Stith, P. M. Burgers, W. D. Heyer, PCNA is required for initiation of recombination-associated DNA synthesis by DNA polymerase delta. Mol Cell 36, 704–713 (2009).

33. R. P. Anand, S. T. Lovett, J. E. Haber, Break-induced DNA replication. Cold Spring Harb Perspect Biol 5, a010397 (2013).

34. A. Atari, H. Jiang, R. A. Greenberg, Mechanisms and genomic implications of break-induced replication. Nat Struct Mol Biol 32, 1871–1882 (2025).

35. L. Liu, A. Malkova, Break-induced replication: unraveling each step. Trends Genet 38, 752–765 (2022).

36. M. A. Wilson et al., Pif1 helicase and Polδ promote recombination-coupled DNA synthesis via bubble migration. Nature 502, 393–396 (2013).

37. O. Buzovetsky et al., Role of the Pif1-PCNA Complex in Pol δ-Dependent Strand Displacement DNA Synthesis and Break-Induced Replication. Cell Rep 21, 1707–1714 (2017).

38. S. Li et al., PIF1 helicase promotes break-induced replication in mammalian cells. EMBO J 40, e104509 (2021).

39. Y. Vasianovich, L. A. Harrington, S. Makovets, Break-induced replication requires DNA damage-induced phosphorylation of Pif1 and leads to telomere lengthening. PLoS Genet 10, e1004679 (2014).

40. K. N. Koc, S. P. Singh, J. L. Stodola, P. M. Burgers, R. Galletto, Pif1 removes a Rap1-dependent barrier to the strand displacement activity of DNA polymerase delta. Nucleic Acids Res 44, 3811–3819 (2016).

41. S. P. Singh, A. Soranno, M. A. Sparks, R. Galletto, Branched unwinding mechanism of the Pif1 family of DNA helicases. Proc Natl Acad Sci U S A 116, 24533–24541 (2019).

42. M. Ortiz-Rodríguez, S. P. Singh, F. J. Cao-García, R. Galletto, B. Ibarra, Regulation of Pfh1 helicase activity by nucleic acid interactions and mitochondrial SSB. Proc Natl Acad Sci U S A 123, e2602528123 (2026).

43. D. Dahan et al., Pif1 is essential for efficient replisome progression through lagging strand G-quadruplex DNA secondary structures. Nucleic Acids Res 46, 11847–11857 (2018).

44. M. Varon et al., Rrm3 and Pif1 division of labor during replication through leading and lagging strand G-quadruplex. Nucleic Acids Res 52, 1753–1762 (2024).

45. N. Saini et al., Migrating bubble during break-induced replication drives conservative DNA synthesis. Nature 502, 389–392 (2013).

46. G. L. Moldovan, B. Pfander, S. Jentsch, PCNA, the maestro of the replication fork. Cell 129, 665–679 (2007).

47. R. Bravo, R. Frank, P. A. Blundell, H. Macdonald-Bravo, Cyclin/PCNA is the auxiliary protein of DNA polymerase-delta. Nature 326, 515–517 (1987).

48. J. M. Gulbis, Z. Kelman, J. Hurwitz, M. O’Donnell, J. Kuriyan, Structure of the C-terminal region of p21(WAF1/CIP1) complexed with human PCNA. Cell 87, 297–306 (1996).

49. E. Warbrick, PCNA binding through a conserved motif. Bioessays 20, 195–199 (1998).

50. N. Mailand, I. Gibbs-Seymour, S. Bekker-Jensen, Regulation of PCNA-protein interactions for genome stability. Nat Rev Mol Cell Biol 14, 269–282 (2013).

51. W. H. Beckwith et al., Destabilized PCNA trimers suppress defective Rfc1 proteins in vivo and in vitro. Biochemistry 37, 3711–3722 (1998).

52. C. N. Ionescu, K. A. Shea, R. Mehra, L. Prundeanu, M. A. McAlear, Monomeric yeast PCNA mutants are defective in interacting with and stimulating the ATPase activity of RFC. Biochemistry 41, 12975–12985 (2002).

53. C. Gaubitz et al., Cryo-EM structures reveal high-resolution mechanism of a DNA polymerase sliding clamp loader. Elife 11 (2022).

54. F. Zheng, R. Georgescu, N. Y. Yao, H. Li, M. E. O’Donnell, Cryo-EM structures reveal that RFC recognizes both the 3’- and 5’-DNA ends to load PCNA onto gaps for DNA repair. Elife 11 (2022).

55. X. Liu, C. Gaubitz, J. Pajak, B. A. Kelch, A second DNA binding site on RFC facilitates clamp loading at gapped or nicked DNA. Elife 11 (2022).

56. X. V. Gomes, S. L. Gary, P. M. Burgers, Overproduction in Escherichia coli and characterization of yeast replication factor C lacking the ligase homology domain. J Biol Chem 275, 14541–14549 (2000).

57. D. Kim et al., DNA skybridge: 3D structure producing a light sheet for high-throughput single-molecule imaging. Nucleic Acids Res 47, e107 (2019).

58. H. S. Kim, S. J. Brill, Rfc4 interacts with Rpa1 and is required for both DNA replication and DNA damage checkpoints in Saccharomyces cerevisiae. Mol Cell Biol 21, 3725–3737 (2001).

59. A. Yuzhakov, Z. Kelman, J. Hurwitz, M. O’Donnell, Multiple competition reactions for RPA order the assembly of the DNA polymerase delta holoenzyme. EMBO J 18, 6189–6199 (1999).

60. V. Ellison, B. Stillman, Biochemical characterization of DNA damage checkpoint complexes: clamp loader and clamp complexes with specificity for 5’ recessed DNA. PLoS Biol 1, E33 (2003).

61. Y. Matsumoto, Molecular mechanism of PCNA-dependent base excision repair. Prog Nucleic Acid Res Mol Biol 68, 129–138 (2001).

62. L. M. Podust, V. N. Podust, J. M. Sogo, U. Hubscher, Mammalian DNA polymerase auxiliary proteins: analysis of replication factor C-catalyzed proliferating cell nuclear antigen loading onto circular double-stranded DNA. Mol Cell Biol 15, 3072–3081 (1995).

63. G. O. Bylund, P. M. Burgers, Replication protein A-directed unloading of PCNA by the Ctf18 cohesion establishment complex. Mol Cell Biol 25, 5445–5455 (2005).

64. Z. Zhuang, B. L. Yoder, P. M. Burgers, S. J. Benkovic, The structure of a ring-opened proliferating cell nuclear antigen-replication factor C complex revealed by fluorescence energy transfer. Proc Natl Acad Sci U S A 103, 2546–2551 (2006).

65. F. Uhlmann, J. Cai, E. Gibbs, M. O’Donnell, J. Hurwitz, Deletion analysis of the large subunit p140 in human replication factor C reveals regions required for complex formation and replication activities. J Biol Chem 272, 10058–10064 (1997).

66. E. Johansson, J. Majka, P. M. Burgers, Structure of DNA polymerase delta from Saccharomyces cerevisiae. J Biol Chem 276, 43824–43828 (2001).

67. G. J. Wuite, S. B. Smith, M. Young, D. Keller, C. Bustamante, Single-molecule studies of the effect of template tension on T7 DNA polymerase activity. Nature 404, 103–106 (2000).

68. F. Cerrón et al., Replicative DNA polymerases promote active displacement of SSB proteins during lagging strand synthesis. Nucleic Acids Res 47, 5723–5734 (2019).

69. G. A. I. Plaza et al., Mechanism of strand displacement DNA synthesis by the coordinated activities of human mitochondrial DNA polymerase and SSB. Nucleic Acids Res 51, 1750–1765 (2023).

70. T. Mondol, J. L. Stodola, R. Galletto, P. M. Burgers, PCNA accelerates the nucleotide incorporation rate by DNA polymerase δ. Nucleic Acids Res 47, 1977–1986 (2019).

