## Supplementary material for "Dissecting the role of PCNA and Pif1 in replication of individual DNA molecules by DNA polymerase δ": Zaher et al_Supplementary Information_20260519.pdf

#### MATERIALS AND METHODS

##### DNA substrates.

The 17,853 bp-long gapped DNA substrate (Lumicks Biotinylated DNA Hybrid SKU: 00027) is nicked at two positions in the same strand so that stretching it at forces above 60 pN generates a central ~5 knt ssDNA gap. The substrate is also asymmetrically labeled with ATTO 647N fluorophore, allowing identification of the 3'-edge of the gap.

Lumicks DNA tethering kit (SKU: 00026) was used to insert a region of interest (ROI) between two ~6 kbp DNA handles, one of which is labeled with the ATTO 647N fluorophore. For one ROI, we constructed a Cy3-labeled short flap substrate ($T_{10}$ Gap-$T_{25}$ Flap in Table S1) by annealing oligonucleotides MZ22: MZ21: MZ23 at a ratio of 1: 1.2: 1.4 in an annealing buffer consisting of 10 mM Tris (pH=8.0), 0.1 mM EDTA (pH=8.0), and 100 mM NaCl. The resulting short flap substrate was ligated to the DNA handles of the Lumicks DNA tethering kit according to manufacturer's suggestions. The reaction was then incubated at room temperature for 1 hr, followed by quenching with 30 mM EDTA and heat inactivation at 65°C for 20 mins. The substrate was used without further processing at a 1000x dilution.

Similarly, other ROIs containing different junctions (Table S1) were annealed and ligated following the same procedure. All oligonucleotides were purchased from Integrated DNA Technologies and are listed in Table S1.

**Table S1.** Oligonucleotide sequences and short DNA substrates used in this study.

| | |
|---|---|
| MZ21 | /5Phos/CAACCACACCGCATATGGTGCACT |
| MZ22 | T/iCy3/TTTTTTTTTTTTTTTTTTTTTTTTCAAGCATTTTATCCGTACTCCTGC |
| MZ23 | /5Phos/ACCAGCAGGAGTACGGATAAAATGCTTGATTTTTTTTTTAGTGCACC ATATGCGGTGTG |
| MZ33 | TCAAGCATTTTATCCGTACTCCTGC |
| MZ34 | /5Phos/ACCAGCAGGAGTACGGATAAAATGCTTGATAGTGCACCATATGCGG TGTG |

| | |
|---|---|
| MZ35 | /5Phos/ACCAGCAGGAGTACGGATAAAATGCTTGATTTTTTTTTTTTTTTTTTTT TTTTTTTTTTAGTGCACCATATGCGGTGTG |
| $T_{10}$ Gap | MZ33 + MZ21 + MZ23 |
| $T_{30}$ Gap | MZ33 + MZ21 + MZ35 |
| $T_{10}$ Gap-$T_{25}$ Flap | MZ22 + MZ21 + MZ23 |
| $T_{1}$ Gap-$T_{25}$ Flap | MZ22 + MZ21 + MZ34 |

**Protein Purifications.**

*Saccharomyces Cerevisiae* ΔN-RFC1 RFC (hereinafter referred to as RFC) was expressed in *E. coli* and purified according to (Gomes, Gary et al. 2000). Unlabeled *Sc*RPA was expressed and purified as described previously for *Hs*RPA (Henricksen, Umbricht et al. 1994). Yeast DNA polymerase δ exonuclease deficient mutant (Pol $\delta^{DV}$, hereinafter referred to as Pol δ) was expressed in yeast and purified according to (Koc, Stodola et al. 2015). MB543-*Hs* and -*Sc*RPA and Alexa Fluor 555-*Ec*SSB were expressed, purified and labeled as described (Roy, Kozlov et al. 2009, Pokhrel, Origanti et al. 2017, Kuppa, Pokhrel et al. 2021). *Sc*Pif1 was purified as in (Singh, Koc et al. 2016) with minor modifications.

Finally, 4C->S-K164C-*Sc*PCNA as reported in (Chen, Ai et al. 2010) was expressed and purified from Rosetta 2 (DE3), with the following modifications. Cultures were grown in TB media at 37°C until $OD_{600}$=0.9, induced with 0.5 mM IPTG and allowed to grow for another 20 hrs at 16°C. Cells were harvested and lysed via sonication in Buffer I (30 mM HEPES-NaOH pH=8.0, 1 mM EDTA, 10 % glycerol, 1 mM DTT) + 600 mM NaCl + 1 mM PMSF + protease inhibitor cocktail tablet. The cleared lysate was treated with 0.1% Polymin P, and the supernatant was dialyzed against buffer containing Buffer I + 300 mM NaCl, diluted 3X to 100mM NaCl and loaded onto a 5 mL HiTrap Heparin HP column (Cytiva) preequilibrated with 5% of Buffer I + 2M NaCl. The flow through from this column was directly loaded onto a 5 mL HiTrap Q HP column (Cytiva) preequilibrated with 5% of Buffer I + 2M NaCl and eluted with a gradient of 5-30% Buffer I + 2M NaCl in 20 CV. Fractions containing the protein were pooled, concentrated, and 10 mM DTT was added, then they were loaded onto a HiPrep 16/60 Sephacryl S200 HR column (Cytiva) equilibrated with Buffer I + 500 mM NaCl and 0.1 mM TCEP in place of 1 mM DTT. Fractions containing the protein were pooled and processed for labeling. Briefly, PCNA-K164C was labeled with AATOM647N - maleimide (AAT Bioquest, 2857) at a 10x molar excess of fluorophore over protein (trimer). The reaction was incubated overnight at 4°C, quenched with 10 mM DTT and excess unreacted fluorophore removed by loading onto a packed Bio-gel P6 media (Bio-Rad) column equilibrated in Buffer I + 500 mM NaCl. The average degree of labeling was ~ 0.7 fluorophore/monomer.

**Single Molecule Experiments.**

*DNA Tethering* – Single molecule experiments were performed using a Lumicks C-Trap instrument that combines dual optical trap tweezers with confocal microscopy and a microfluidic system. Experiments were controlled and data acquired using Bluelake software (versions 2.7.2-2.7.5). A flow cell containing 5 barrierless channels was used to house the experimental setup (Supplementary Figure S1A). The first three channels were used to tether DNA to ~4.4 μm streptavidin-coated polystyrene beads (Spherotech NC0823438). Beads trapped in channel 1 (1xPBS) were moved into channel 2 (1xPBS) containing the biotinylated DNA substrate to make a DNA tether whose formation was verified by a short force-extension curve (from 0 pN to 20 pN). To make gapped DNA substrates, the single DNA tether was moved into channel 3 (0.1x PBS), stretched to ~8.2 μm (~60 pN), and then rapidly relaxed to 2 μm. Flow was turned off at this time, and the successful generation of the gapped DNA was verified by force-extension curve. For all other substrates, after a DNA tether is formed in channel 2, the DNA tether was moved to channel 3 (0.1x PBS) without flow and the presence of a single tether was verified by a force-extension curve. Confocal imaging was performed using a red (638 nm) and/or a green laser (532 nm) at low power (1 to 5%) to identify the position of the ATTO 647N or Cy3 fluorophore with respect to the attached tether. Kymographs were generated by scanning a line drawn between the center of the trapped beads. After identification of its position, the ATTO 647N fluorophore on the DNA tether was then purposefully photobleached by increasing the red laser power to >20%. This photobleaching step was introduced to avoid interference with the AATOM 647N fluorophore on PCNA. The DNA tethers made in channel 3 were then moved to channels 4 and 5 to bind the different proteins used in different experiments, as described below.

*PCNA Loading* − Gapped DNA tethers formed in channel 3 were moved to channel 4 containing 0.5 nM MB543-RPA and after ~1 min incubation to allow binding of RPA to the ssDNA gap, flow was turned off, and the RPA-coated gap was imaged with the green laser. The RPA-coated gapped substrate was then moved to channel 5 containing 5-10 nM AATOM 647N-PCNA, 5-10 nM RFC, 1 mM ATP, with or without 10 nM Pol $\delta^{DV}$, and after ~1-2 min incubation, flow was turned off and the whole assembly was moved out of channel 5 into an upstream location along channel 4 (see location ** in Supplementary Figure S1A) and imaged with both green and red lasers. Both channels 4 and 5 contained Reaction Buffer: 50 mM HEPES-NaOH 7.5, 35 mM potassium glutamate, 7 mM MgCl2, 1 mg/mL BSA, 1 mM DTT, 0.4 % Dextrose, 4 mM Trolox, 0.4 mg/mL Catalase and 1 mg/mL glucose oxidase. Loading of PCNA onto the doubly nicked DNA and other DNA substrates (Table S1) followed similar procedure but without MB543-RPA incubation, PCNA in channel 4 and imaging occurred in channel 5 in the presence of other proteins (Pif1, Pol δ, unlabeled RPA).

*Gap-filling synthesis* − 0.5 nM MB543-RPA and 5-10 nM AATOM 647N-PCNA along with 5-10 nM RFC and 1 mM ATP were placed in channel 4 while 30 nM Pol $\delta^{DV}$ and dNTPs (500 μM each) were placed in channel 5, with both channels containing Reaction Buffer. Hence, the gapped substrate was incubated in channel 4 to coat the gap with RPA and load PCNA, then moved in the absence of flow to channel 5 for gap-filling synthesis and imaged with both green and red lasers.

*Strand displacement synthesis* − 5-10 nM AATOM 647N-PCNA, 5-10 nM RFC and 1 mM ATP were placed in channel 4, while 6 (+) or 60 nM (++) Pol δ, dNTPs (500 μM each), 9 (+) or 90 nM (++) Pif1, 1 mM ATP and 20 nM unlabeled RPA were placed in channel 5. Both channels contained

Reaction Buffer. $T_{10}$ gap-$T_{25}$ flap generated in channel 3 was moved to channel 4 to load PCNA, then moved to channel 5 where flow was turned off and imaged with green and red lasers.

**Data Analysis.**

All data analyses were performed using Python scripts implemented with the help of chatgpt 5.3-5.5.

*Gap Length* – To estimate the length of gaps coated by labeled RPA (Supplementary Figure S1B and S1C), an intensity-based approach was utilized to examine each kymograph's green-channel. For each kymograph, the intensity was averaged along the time-axis to yield a 1D distance-intensity profile. The profile was smoothed using a Gaussian filter ($\sigma$ = 2 pixels) and an intensity threshold defined as 20% of the maximum smoothed intensity. The largest contiguous region per kymograph with signal above this threshold was measured in $\mu$m.

*Lifetimes* – For lifetime analysis (Figure 1C-E, Figure 3B and 3C, Supplementary Figures S2, S3D, and S3E), first, the edges of the gap, representing the 3'- and 5'-end, were detected using the green channel, then lifetimes were extracted at these edges using the red channel. A rectangular region of interest (ROI) that excluded the beads was selected for each kymograph and the edges of the green signal detected using a threshold-based approach. Briefly, the image was smoothed, the background estimated as the 5th percentile of the smoothed intensity distribution, and a threshold based on the maximum intensity applied. Lower and upper edges of the gap were defined as the largest contiguous segments of pixels above the threshold and their traces overlayed on the kymograph. Each edge was tagged as "3'-edge" or "5'-edge" according to the position of the ATTO 647N fluorophore on the DNA tether. Once the edges were detected, red-channel intensity profiles were extracted, and at each time point, intensity values were averaged over a fixed-width band of 4 pixels in the distance axis. This allowed for minimizing pixel-level noise and small edge position uncertainties while capturing local intensity fluctuations. Intensity-based lifetimes were then calculated as two states "high" or "low" along each edge based on a threshold defined as a fixed fraction (0.1) of the maximum intensity of that edge. Frames that are above this threshold were identified and the duration calculated and registered as a lifetime. For 0.1s and 0.2s binning, the red intensity in the width of these bins was averaged and a threshold was applied as above. Lifetimes were then sorted and counted according to different criteria based on the tag of the edge and their duration. Lifetimes in Figures 2D, E and 4B, C were estimated by visually inspecting the kymographs with stable red signal (>2s) at the particular site. These lifetimes were filtered with the mean $\pm$ 2x standard deviation.

*Diffusion Coefficient* – To estimate diffusion coefficients (Supplementary Figure S3), ROIs were selected on each kymograph's red channel to choose areas where PCNA was diffusing rather than stationary. To track PCNA trajectories, Lumicks Pylake's track_greedy algorithm was used, and the detected tracks were refined using Pylake's Gaussian localization. Diffusion coefficients were then estimated using Pylake's covariance-based estimator (CVE).

*Time-Averaged Intensity Profiles of PCNA* – For PCNA "localization" on the doubly nicked substrate, time-averaged intensity profiles were computed from the red channel (Figure 2C). To ensure equal temporal weighting, the kymographs were sliced temporally to the shortest kymograph duration included in the dataset. Time-averaged 1D intensity profiles were smoothed by a Gaussian filtering, the bead-DNA boundaries detected and excluded, and distances in micrometers rescaled to base-pair by normalizing to the full DNA length (17,853 bp). To plot all intensity profiles on a common base-pair axis, individual profiles were interpolated and then summed to generate a composite summed intensity profile. The polarity of the tethers was confirmed using the position of the ATTO 647N fluorophore on the DNA, and kymographs were re-oriented accordingly. For DNA contour length (L) and (σ) (Supplementary Figure S4), L was determined as above. As for σ, the position of ATTO 647N of the DNA was localized by taking the maximum of the time-averaged intensity profile within the DNA region and was fit locally with a Gaussian function with standard deviation (σ).

*Gap-Filling Synthesis* – To analyze gap-filling synthesis, two methods were used; tracking RPA displacement and following bead-to-bead distance change. In the first, the edges of the RPA-coated gap were detected from the green channel as described above for lifetime analysis. Edges were tagged as "moving" or "stationary" corresponding to "3'-edge" and "5'-edge", respectively, and were analyzed separately; 1) "moving" edge resulting from loss of green signal due to RPA displacement upon DNA synthesis, and 2) "stationary" edge with stable RPA signal because no DNA synthesis occurs from the 5'-end. For the bead-to-bead distance change, distance 1 versus time was extracted from .H5 files.

*Strand Displacement Synthesis* – Directional movement of the Cy3 fluorophore on the 5'-flap of $T_{10}$ gap-$T_{25}$ flap substrate was taken as a proxy for DNA synthesis. To track the Cy3 signal, ROIs were manually selected to exclude bead edges, and the green channel signal was tracked using Pylake's track_greedy algorithm and position refined using Gaussian localization. Tracks were converted to time-position coordinates and points that deviates beyond 0.7 standard deviations from the local mean were filtered out using a rolling window.

*Multi segment fitting* – The digitized traces corresponding to either the edges of the RPA gap, the bead-to-bead distance, or the Cy3 signal of the 5'-flap were segmented into piecewise linear fragments. First, the traces were smoothed, and candidate inflection points were detected using the second temporal derivative. Then, each segment was fitted using least-squares regression. To choose an optimal number of segments, a cost matrix was built, and Bayesian Information Criterion (BIC) was minimized. Furthermore, a likelihood-ratio F-test was used to compare nested models of increasing complexity. Additionally, minimum segment duration and a minimum temporal separation between consecutive breakpoints constrained the segmentation. Operationally, we also used a visual inspection approach to conservatively select the minimum number of segments. Finally, DNA synthesis rates were calculated from the slopes of the linear fits, and track lengths of newly synthesized DNA were calculated as the differences in distance between the start and end of the segments. For analysis of gap-filling using RPA displacement, we used the average slope of the "stationary" edge as a threshold to filter out from the "moving" edge analysis segments with slopes below that threshold. These segments were considered as segments representing pauses in DNA synthesis. As for analysis using bead-to-bead distance, only segments with negative slopes were considered. They were further filtered by the average slope of segments of the DNA alone kymographs and kymographs that showed no DNA synthesis both with RPA displacement and bead-to-bead distance change. For analysis using the Cy3 signal of the 5'-flap, segments were filtered using the slope of segments corresponding to the flapped substrate in the absence of any protein.

Conversion from μm to nt was performed using the estimated gap length in μm corresponding to 5005 nts for calculating rates and track lengths by tracking RPA displacement. For bead-to-bead distance change, the conversion followed the FD curves (Supplementary Figure S5D). Finally, for strand displacement synthesis, the conversion rate followed the full length of the DNA from bead-edge to bead-edge (3.4 μm) corresponding to 12596 nt total. The rates and track lengths (Figure 3F and 4E) were further filtered with the mean ± 1.5x standard deviation.

#### Supplementary References:


Chen, J., Y. Ai, J. Wang, L. Haracska and Z. Zhuang (2010). "Chemically ubiquitylated PCNA as a probe for eukaryotic translesion DNA synthesis." Nat Chem Biol **6**(4): 270-272.

Gomes, X. V., S. L. Gary and P. M. Burgers (2000). "Overproduction in Escherichia coli and characterization of yeast replication factor C lacking the ligase homology domain." J Biol Chem **275**(19): 14541-14549.

Henricksen, L. A., C. B. Umbricht and M. S. Wold (1994). "Recombinant replication protein A: expression, complex formation, and functional characterization." J Biol Chem **269**(15): 11121-11132.

Koc, K. N., J. L. Stodola, P. M. Burgers and R. Galletto (2015). "Regulation of yeast DNA polymerase δ-mediated strand displacement synthesis by 5'-flaps." Nucleic Acids Res **43**(8): 4179-4190.

Kuppa, S., N. Pokhrel, E. Corless, S. Origanti and E. Antony (2021). "Generation of Fluorescent Versions of Saccharomyces cerevisiae RPA to Study the Conformational Dynamics of Its ssDNA-Binding Domains." Methods Mol Biol **2281**: 151-168.

Pokhrel, N., S. Origanti, E. P. Davenport, D. Gandhi, K. Kaniecki, R. A. Mehl, E. C. Greene, C. Dockendorff and E. Antony (2017). "Monitoring Replication Protein A (RPA) dynamics in homologous recombination through site-specific incorporation of non-canonical amino acids." Nucleic Acids Res **45**(16): 9413-9426.

Roy, R., A. G. Kozlov, T. M. Lohman and T. Ha (2009). "SSB protein diffusion on single-stranded DNA stimulates RecA filament formation." Nature **461**(7267): 1092-1097.

Singh, S. P., K. N. Koc, J. L. Stodola and R. Galletto (2016). "A Monomer of Pif1 Unwinds Double-Stranded DNA and It Is Regulated by the Nature of the Non-Translocating Strand at the 3'-End." J Mol Biol **428**(6): 1053-1067.

#### SUPPLEMENTARY FIGURE S1

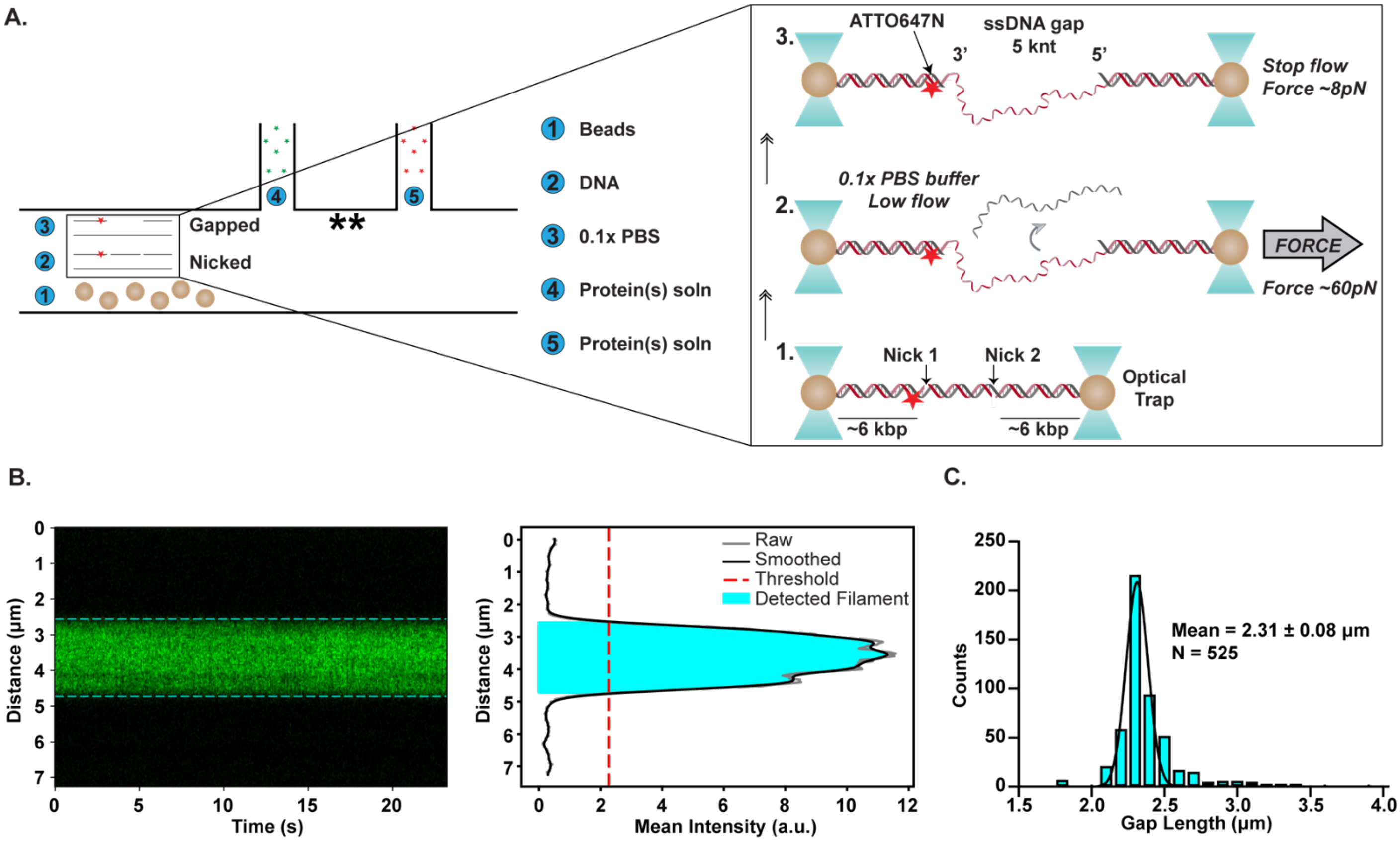


**Supplementary Figure S1. Experimental design. A)** A schematic illustrating the microfluidic flow cell with 5 barrierless channels. Channel 1 contained the beads that were optically trapped and then moved to channel 2 where a tether of nicked DNA was established. The tethered DNA was then moved to channel 3 to construct the gapped substrate, as depicted in the inset to the right. The tether was moved to channel 3 where a low flow of 0.1x PBS buffer was maintained as the DNA was stretched to a ~60 pN force and an 8.2 μm distance, and then rapidly relaxed. The generation of 5 knt ssDNA gap was confirmed with force-extension curves. The flow was turned off, and the DNA was stretched to ~8 pN (passive mode) or clamped at the desired force and then moved into channels 4 and 5 to bind to proteins and image. (**) denotes the position where loaded PCNA was imaged when PCNA was in channel 5. **B)** A kymograph showing an RPA-coated gapped substrate (green) with the edges denoted by dashed cyan lines (Left). The intensity of the green signal was averaged over the time axis, smoothed and a threshold was applied to detect the length of the RPA-coated gap along the distance axis (Right). **C)** A bar chart showing the Gaussian fitted frequency distribution of gap lengths of 525 kymographs. The reported values are the mean ± standard deviation.

#### SUPPLEMENTARY FIGURE S2

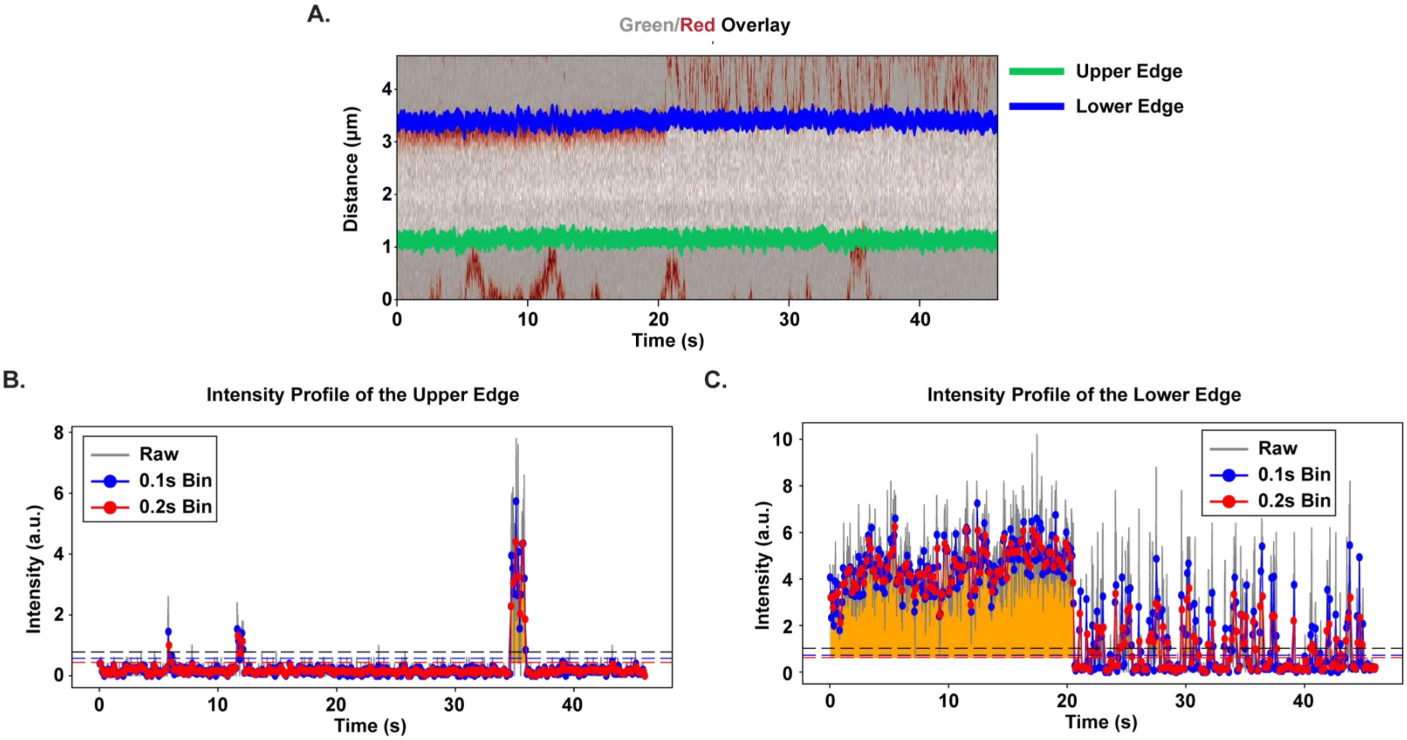


**Supplementary Figure S2. Lifetime analysis**. **A)** A kymograph of RPA-coated gap substrate (green channel shown in gray scale) and loaded PCNA (in red) with the upper (green) and lower (blue) edges of the gap detected by thresholding and defined as a band of 4-pixel width. **B, C)** Intensity profiles of the red channel (i.e. PCNA) along the upper **(B)** and lower **(C)** edges detected in (A) were averaged over the 4-pixel width and plotted against time. The signal was then binned with either 0.1s or 0.2s bins to smooth the data and the lifetimes measured as described in Materials and Methods. Lifetimes in Figure 1D and Figure 3C are binned at 0.2s.

SUPPLEMENTARY FIGURE S3

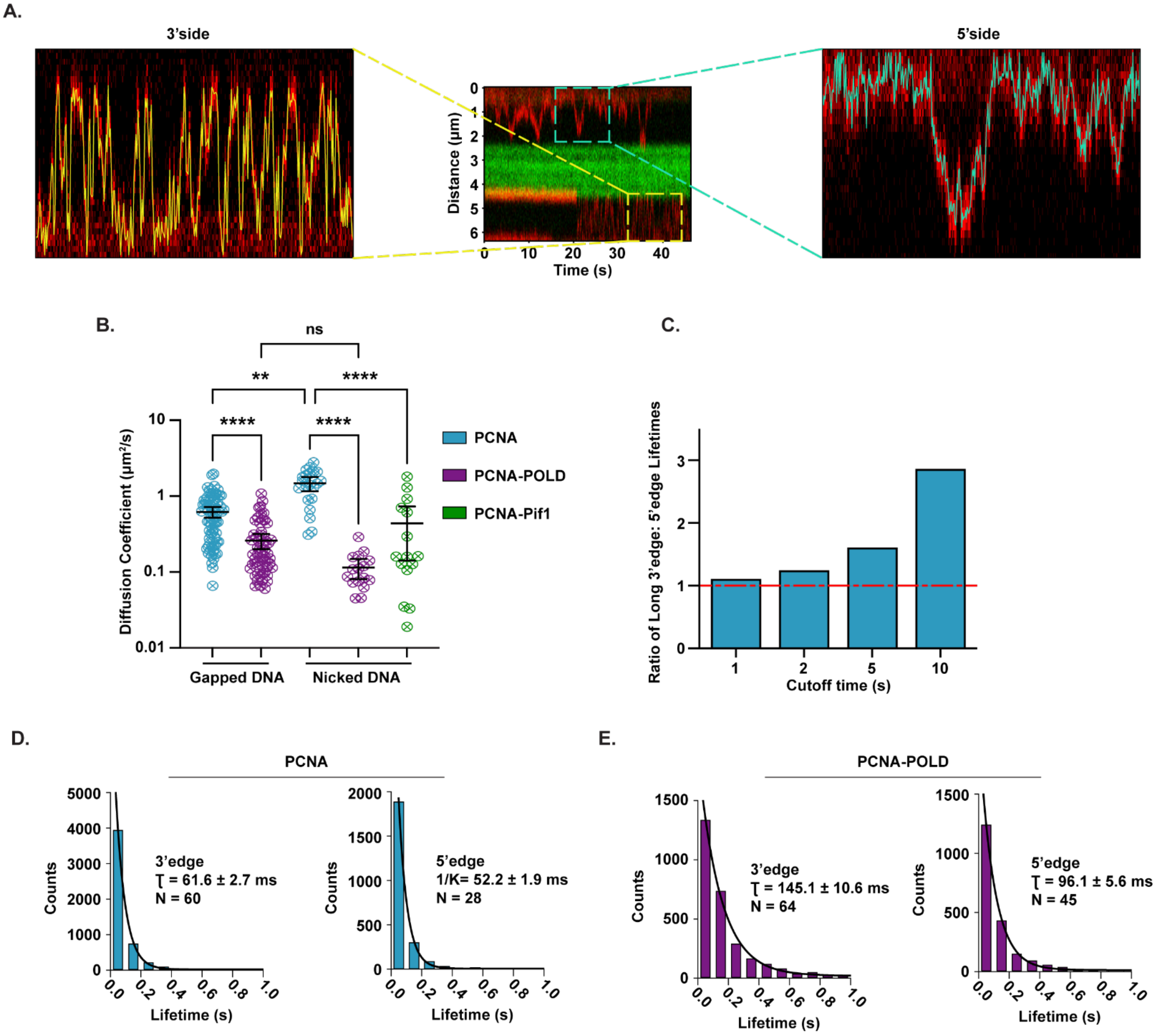


**Supplementary Figure S3. Analysis of the 1D diffusing mode of PCNA and its short-lived states at the edges. A)** Kymograph from Figure 1A showing diffusive properties of loaded PCNA on the 3' side (Left) and 5' side (Right). **B)** Diffusion coefficients of loaded PCNA in the presence and absence of Polδ on gapped or nicked DNA substrates. Error bars represent the mean ± 95% confidence interval and statistical significance was determined by Kruskal-Wallis test. **C)** Bar chart of the ratio of the 3'-to 5'-edge long-lived lifetimes, calculated at different cutoff times. A ratio above 1 indicates that, independent of the cutoff chosen, the long-lived states dominate at the 3'- versus 5'-edge. **D, E)** Exponentially fitted frequency distributions of short-lived (<2s) lifetimes (unbinned raw data) at the 3'-edge or 5'-edge of a gap in the absence **(D)** or presence of Polδ **(E).** The reported values are the mean ± 95% confidence interval.

#### SUPPLEMENTARY FIGURE S4

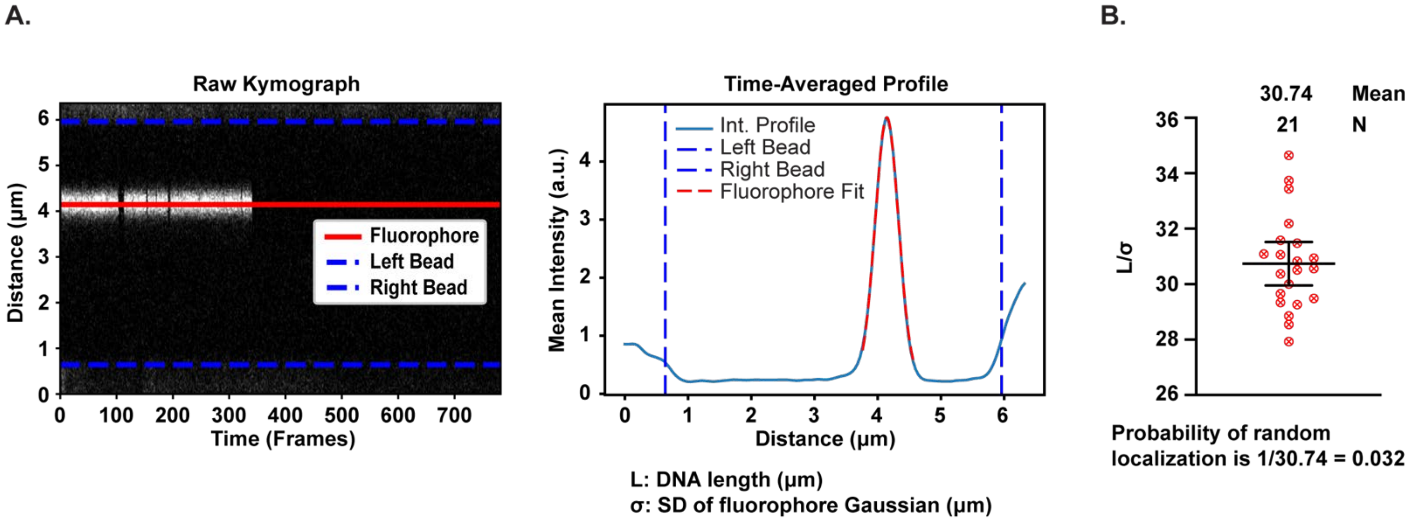


**Supplementary Figure S4. Probability of a PCNA event to be positioned randomly on a nicked DNA substrate. A)** Left: a kymograph showing the position of the ATTO 647N fluorophore on a nicked DNA substrate (red line) between two beads, the edges of which were detected and denoted by dashed blue lines. Right: time-averaged intensity profile of the fluorophore shows the edges of the beads (dashed blue lines) and Gaussian fit of the fluorophore intensity. From this intensity profile, two parameters were calculated; L: the length of the DNA from bead-to-bead edge and $\sigma$: the standard deviation of the fluorophore Gaussian denoting the spread of the fluorophore. **B)** A plot of $L/\sigma$ denoting the minimum number of fluorophores that can be accommodated on the length of the DNA given their spread which has a mean of 30.74. Therefore, the probability of a fluorophore being at a random position along the DNA is at a maximum of 0.032 or 3.2%. Error bars represent the mean $\pm$ 95% confidence interval

### SUPPLEMENTARY FIGURE S5

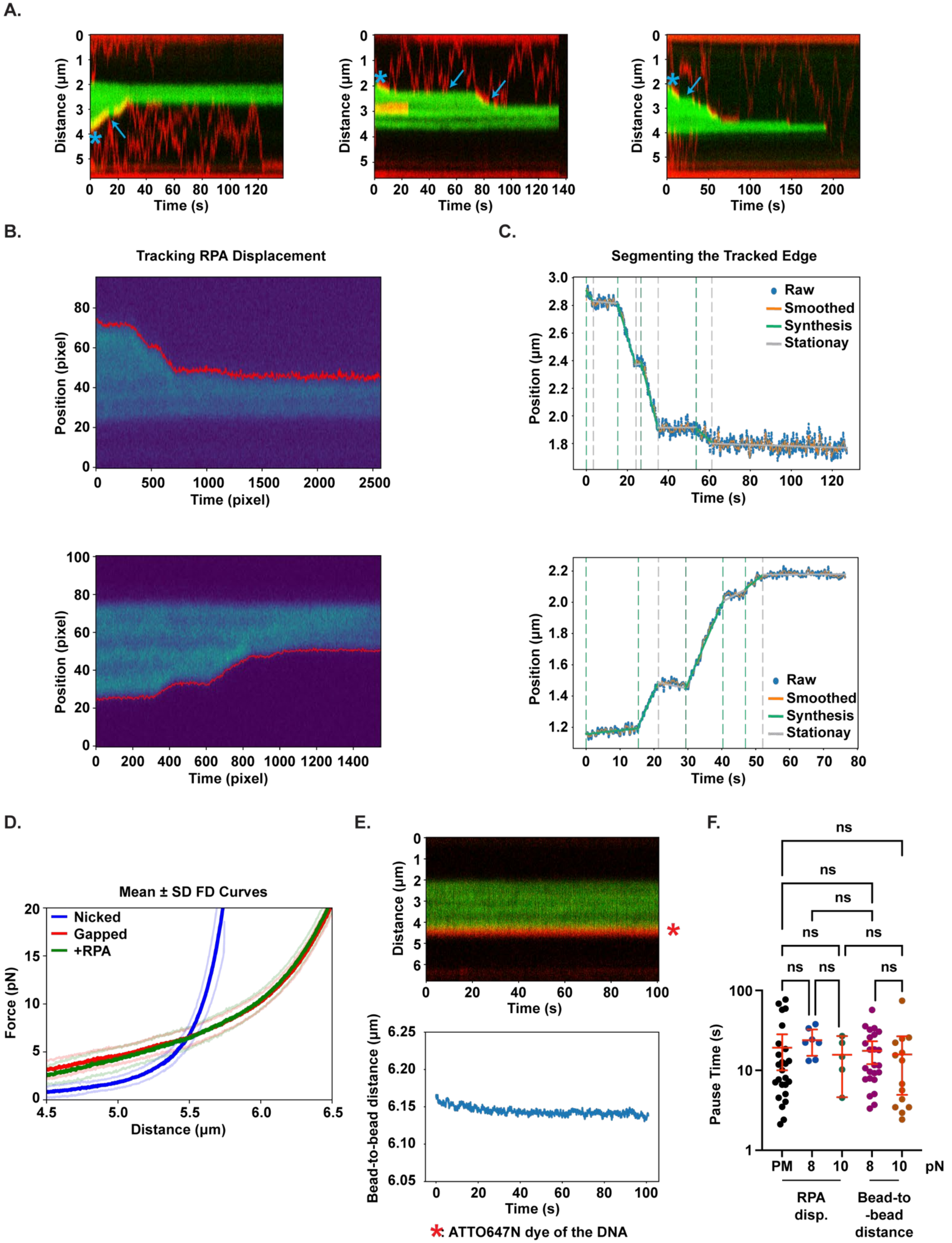

**Supplementary Figure S5. Gap-filling synthesis rate, processivity and pause-time calculation. A)** Additional kymographs following gap-filling synthesis in the presence of Polδ and dNTPs in passive mode, as shown in Figure 3D. **B)** To determine rate, processivity and pause times by tracking RPA displacement, the edge where RPA is being displaced was tracked by thresholding as described in Materials and Methods. **C)** The tracked edge was then plotted, smoothed and segmented and the segments were classified as synthesis or stationary by comparing their slopes to the slopes of the segments of the tracked and segmented non-displaced RPA edge. **D)** Mean (solid color) ± standard deviation (faded color) of Force-Distance (FD) curves of nicked DNA (blue), gapped DNA (red) and RPA-coated gapped DNA (green). **E)** Top: a kymograph showing RPA-coated gapped substrate in the presence of Polδ and dNTPs but in the absence of loaded PCNA. The stationary red signal denotes the ATTO 647N fluorophore of the DNA substrate and thus the 3' edge of the gap. No significant RPA displacement was witnessed. Bottom: bead-to-bead distance as a function of time of the same kymograph showing no significant change. **F)** Gap-filling DNA synthesis pause times under different conditions; passive mode (PM) or under 8 or 10 pN force, and as calculated by different methods; following RPA displacement or the change in bead-to-bead distance. Error bars represent the mean ± 95% confidence interval and statistical significance was determined using a Kruskal-Wallis test.

#### SUPPLEMENTARY FIGURE S6

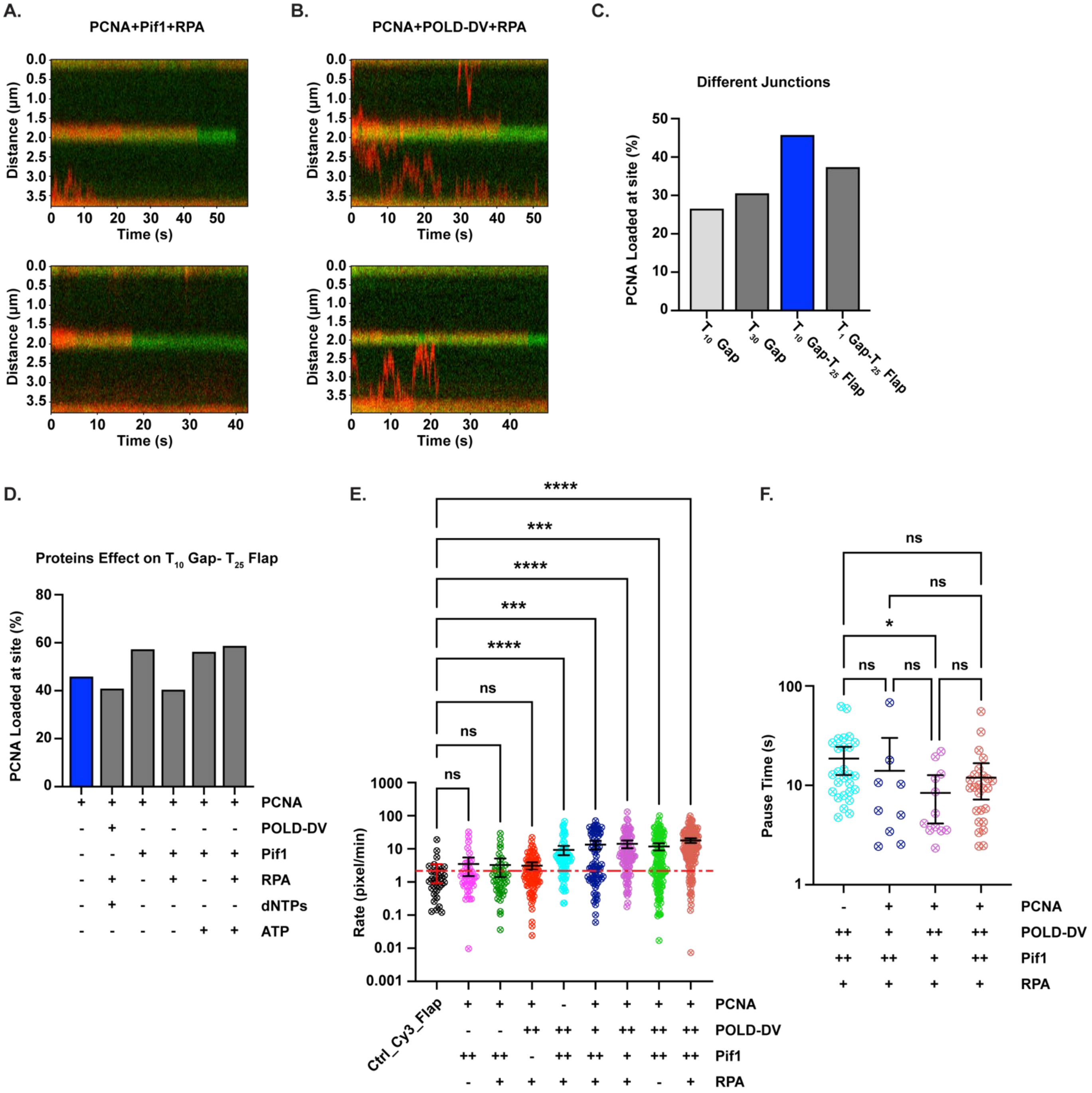


**Supplementary Figure S6. Strand displacement DNA synthesis requires the activity of Polδ and the Pif1 helicase**. **A, B)** Kymographs following the signal of the Cy3-labeled 5'flap (green) of the $T_{10}$ gap-$T_{25}$ flap substrate along with loaded PCNA (red) in the presence of Pif1, ATP and unlabeled RPA **(A)** or in the presence of Polδ, dNTPs, and unlabeled RPA **(B)**. **C)** Percentage of loaded PCNA at a $T_{10}$ gap, $T_{30}$ gap, $T_{10}$ gap-$T_{25}$ flap or $T_1$ gap-$T_{25}$ flap. **D)** Percentage of loaded PCNA at a $T_{10}$ gap with $T_{25}$ 5'-flap in the presence of the indicated combination of proteins and nucleotides. **E, F)** Strand-displacement synthesis rate and pause times in the presence of different combinations of proteins. The dashed red line in E) denotes the mean of the rates of the movement of the Cy3-labeled 5'-flap in the same experimental buffer in the absence of any proteins. It is the baseline with which the rates of the other experiments were filtered (Figure 4E). Error bars in E) and F) represent the mean ± 95% confidence interval and statistical significance was determined using a Kruskal-Wallis test.
